# Dynamic context-based updating of object representations in visual cortex

**DOI:** 10.1101/2025.02.05.636616

**Authors:** Giacomo Aldegheri, Surya Gayet, Marius V. Peelen

## Abstract

Objects in real-world scenes are often poorly or partially visible, for example because they are occluded or appear in the periphery. An additional challenge of real-world vision is that it is dynamic, causing the appearance of objects (e.g., their size and orientation) to change as we move. Importantly, however, these changes are predictable from the 3D structure of the surrounding scene. In two fMRI studies, we find that visual cortex dynamically updates object representations using this predictive contextual information. Firstly, visual cortical representations of objects were enhanced when they rotated congruently (versus incongruently) with the surrounding scene. Secondly, the inferred orientation of the object could be decoded from visual cortex activity even when the object was fully occluded. These findings indicate that predictive processes in visual cortex follow the geometric structure of the environment, providing a mechanism to support object perception in dynamic natural vision.

**Teaser:** Objects are mentally rotated together with the changing viewpoint on a scene, affecting their representation in visual cortex.

## Introduction

Real-world vision is inherently inferential (*1–3*). For example, when part of a scene is occluded, we use contextual information to infer the occluded parts (*4*). Recent research has shown that such perceptual inferences activate regions of visual cortex that are also activated during stimulus-driven perception. For example, neuroimaging studies in humans (*5–7*), and electrophysiological recordings in non-human primates (*8*) and rodents (*9*), revealed that patterns of neural activity in early visual cortex (EVC) contained information about occluded parts of scenes. Similarly, neuroimaging studies showed that scene context modulated late visual cortex (LVC) representations of degraded and poorly visible objects, such that these representations became more similar to the representations of fully visible objects (*10*, *11*). These studies show that perceptual inferences based on (static) scene context do not only affect higher-level decisional stages (*12*) but also modulate and activate visual cortex representations, thereby shaping our perceptual experience (*13*, *14*).

Perceptual inferences in the real world, however, are not only based on static context. As we move, our view of a scene - and the objects within that scene - changes. These changes depend on geometric constraints such as the way a 3D rotation results in a 2D image change on the retina. Importantly, inanimate objects (e.g., a bed) usually remain stable relative to the scene background (e.g., a room). This allows for predicting the appearance of objects from new viewpoints based solely on viewing the scene background. In a recent behavioral study, we found that temporarily occluded objects placed in scenes were indeed automatically mentally rotated together with the changing viewpoint of the surrounding scene (*15*). Specifically, participants performed better on a challenging change discrimination task on the visual object, when the object re-appeared in an orientation that was consistent with the (now rotated) background scene. Because the amount of scene rotation was unpredictable in that study, the new viewpoint of the object could only be inferred from the new viewpoint of the scene, and not through continuous mental rotation of the object alone. This finding provides evidence that predictions of 3D object rotations can occur automatically, as a product of contextual information (in a subsequent study, we found this to occur for translation as well as rotation (*16*)). To our knowledge, it is unknown whether such dynamic context predictions modulate and/or activate visual cortex activity in the way that static context predictions do. Are visual object representations (i.e., predictions about the 2D appearance of an object) dynamically updated to account for changes in scene viewpoint? If so, this would entail that changes in the 3D scene context are used to generate predictions about the (novel) appearance of an object within that scene, potentially supporting object perception in dynamic, 3D, real-world environments.

Here, we used fMRI to address this question. In Experiment 1, we tested for modulatory effects of dynamic context predictions in visual cortex. Specifically, we hypothesized that visual cortex representations are enhanced when objects re-appear in a viewpoint that is congruent rather than incongruent with the (new) scene viewpoint. Note that we adopt an operational definition of “object representations”, referring specifically to the proximal shape of an object’s projection on the retina — “wide” versus “narrow” — as decoded via multivariate pattern analysis. In Experiment 2, we went one step further and tested whether dynamic context predictions of object appearance not only modulate but also directly activate visual cortex. That is, we tested whether information about the new object orientation (derived from the scene viewpoint) would be present in visual cortex, even when the object itself is still occluded and thus fully invisible. If so, this would provide an important generalization of studies investigating static context predictions (*5*, *6*, *8*) or predictions involving highly simplified stimuli (*17*, *18*) to the complexity of real-world environments.

In both fMRI studies, we focused on two regions of interest (ROIs) within the visual cortex: early visual cortex (EVC; Brodmann areas 17 and 18), given its known role in the completion of partially visible scenes (*5*, *6*, *8*), and late visual cortex (LVC; Brodmann areas 19 and 37), which has been implicated in context-driven inference of object properties (*10*, *11*, *19*). In Experiment 1, we decoded, from activity patterns in these two ROIs, the proximal (i.e., 2D) shape of objects that, after an occlusion period, reappeared oriented congruently or incongruently with the rotation of the surrounding scene (**Figure 1A**). Critically, the initial viewpoint and amount of rotation were chosen such that objects could reappear either in a ‘wide’ or ‘narrow’ projection on the 2D image plane (e.g., a bed viewed from the side, versus the tail end). We found that representations of congruent objects, relative to incongruent objects, were enhanced in EVC, as demonstrated by better discriminability of multivariate activity patterns (i.e., ‘wide’ versus ‘narrow’ decoding). This enhancement was accompanied by an overall lower activation at the whole-brain level, in line with effects of other forms of expectations in visual cortex (*20*, *21*). In Experiment 2, we directly decoded the proximal shape of these same objects, but now during the period of occlusion (while no object was visible on the screen), to determine whether object representations were updated coherently with the rotation of the scene context. Here, we found that proximal object shape could be reliably decoded in LVC, providing evidence for purely top-down driven activity reflecting the predicted object orientation, solely derived from the new scene viewpoint.

Together, these results indicate that scene completion in human visual cortex generalizes to the prediction of object appearance across viewpoint changes in 3D scenes, providing a potential mechanism for efficiently processing partially visible scenes in dynamic real-world environments.

**Figure 1.**
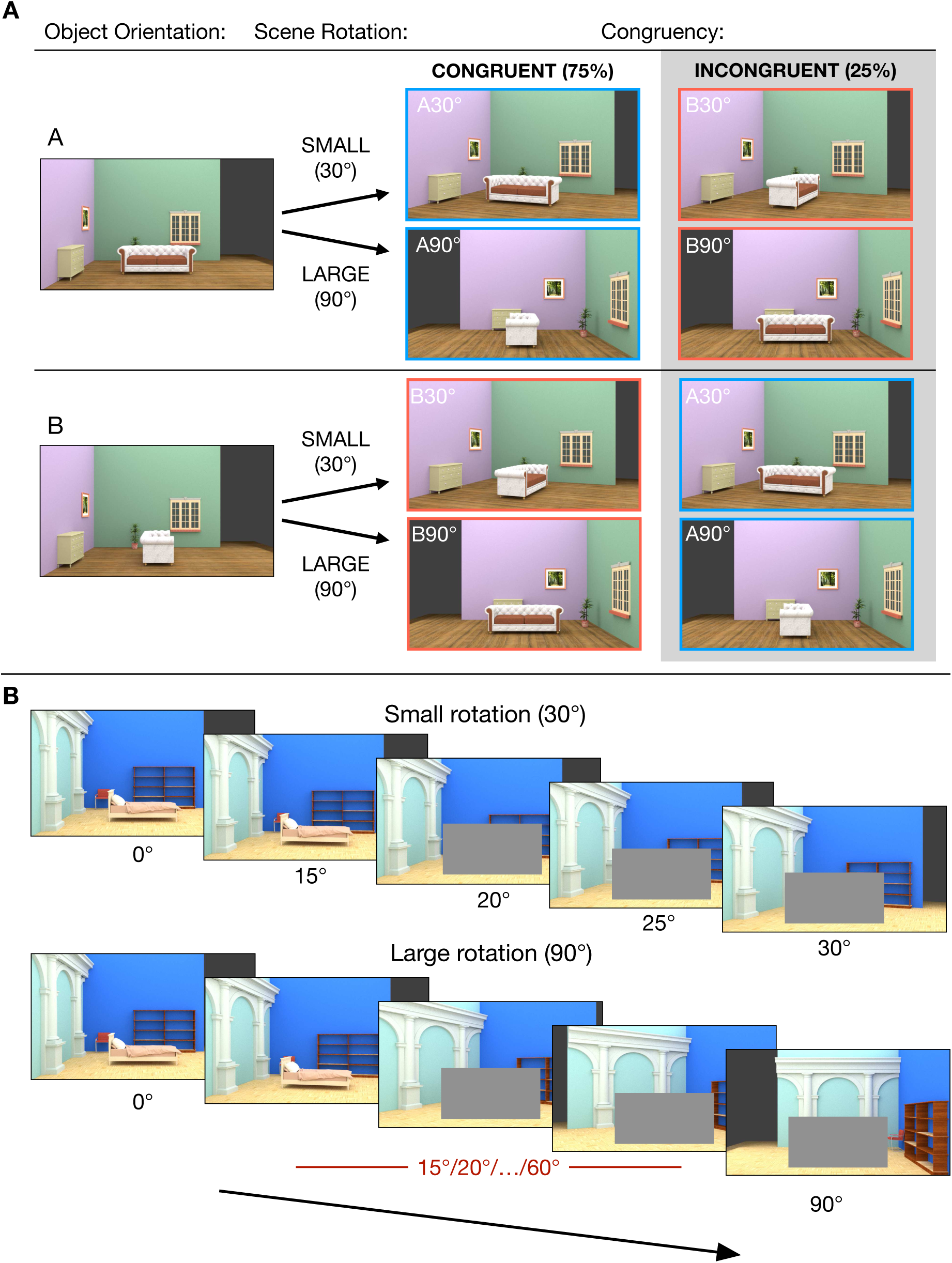
Experimental design of Experiment 1. **(A)** Outline of the experimental design. The stimuli were images of rooms containing a central object, which could be shown at one of two possible orthogonal orientations (labeled A and B) relative to the room. The room could undergo two different total amounts of rotation – small (30°) and large (90°). After the room’s rotation, the object could be either in a Congruent view (with the same orientation relative to the room as at the beginning of the trial) or in an Incongruent view (with the other possible orientation – B if the initial orientation was A or vice versa). **(B)** Examples of the full rotation sequence for a small and large rotation. The rotation was shown in discrete steps, and the object was fully occluded after the first two rotation steps until the whole rotation was complete. The intermediate scene orientations (including the last visible object orientation and the occluded orientations) were fixed (15°-20°-25°) for the 30° rotation trials, and randomly sampled from the set{15°, 20°, 25°, 30°, …, 55°, 60°} for the 90° rotation trials, always shown in increasing order.

**Figure 2.**
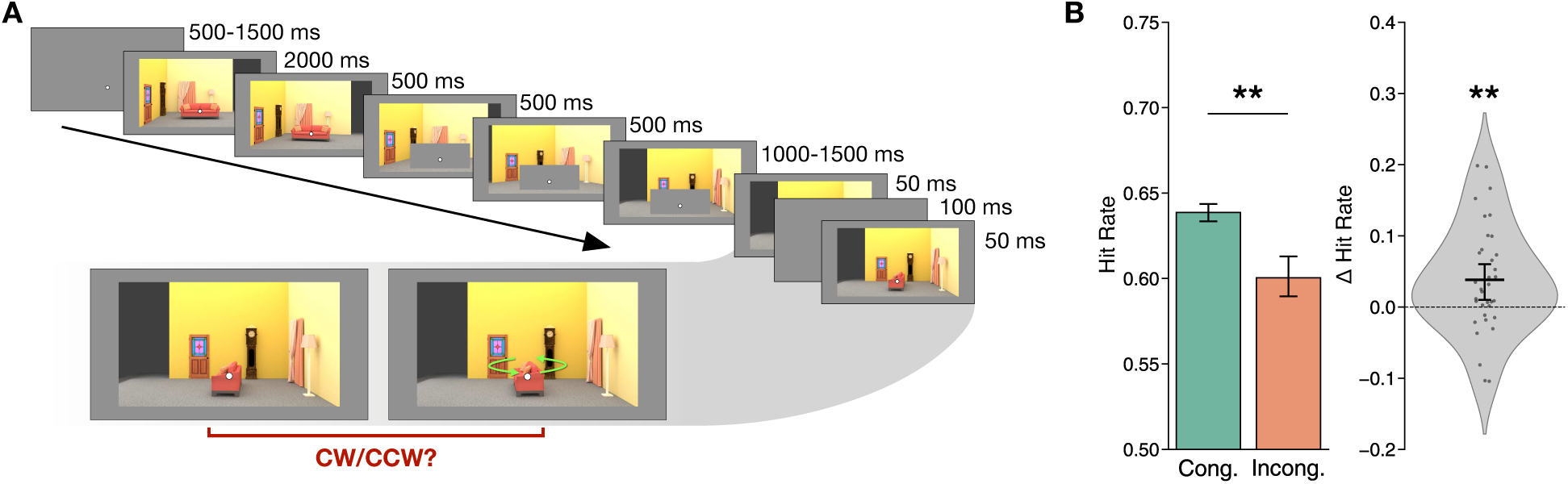
Trial sequence and behavioral results of Experiment 1. **(A)** Temporal outline of a trial. After the rotation was complete, the occluder would disappear, revealing the object in either the Congruent or Incongruent view. The object would be briefly flashed (50 ms) twice, in two slightly different orientations. Participants had to determine whether the second orientation was clockwise or counterclockwise relative to the first. This task was fully orthogonal to the congruency of the object’s orientation, thus allowing us to test whether participants would automatically predict the orientation of the object from the (current) viewpoint on the surrounding scene. **(B)** Mean (and SEM) accuracy on the behavioral task for Congruent and Incongruent trials (left) and distribution of differences in accuracy (Congruent minus Incongruent) across participants. Participants were more accurate when the object’s final view was Congruent. ** P < 0.01

## Results

### Experimental design

In both fMRI experiments, participants viewed realistic indoor scenes (rooms) featuring a central object (a bed or couch) oriented in one of two possible angles relative to the scene (**Figure 1A**). On each trial, the scene would start rotating around the vertical axis in discrete snapshots, causing a change in scene viewpoint **(Figure 1B**). During the first two snapshots the object was fully visible, so that participants could learn how the object was positioned within the room. During the subsequent three snapshots the object was occluded, so that participants would only see the rotating room. In the last snapshot, which occurred on every trial of Experiment 1, the occluder was removed, so that the object became visible again (**Figure 2A**). Critically, the object reappeared in an orientation that was either congruent or incongruent with its original positioning within the room (**Figure 1A**). The total amount of rotation (from initial to final viewpoint) was either 30° or 90°, with each rotation amount occurring on half of the trials within each condition. The amount of rotation on a given trial remained unknown before the object was occluded. Therefore, the new orientation of the object could only be inferred from the new orientation of the room. Importantly, the exact same stimuli (initial and final viewpoints) were used for trials with congruently and incongruently rotated objects. Thus, whether an object was rotated congruently or incongruently could only be inferred through dynamic updating of the object orientation, based on the changing viewpoint on the scene.

Another key aspect of the design is that the two initial object orientations and the two scene rotation angles were chosen to result in two categorically distinct proximal object shapes in the final snapshot: either a *wide* or a *narrow* shape (i.e., the object evoked a wide or narrow projection on the 2D image plane). This was done to maximize the power of the multivariate decoding analyses, discriminating between patterns of activity evoked by wide versus narrow shapes.

### Enhanced representations of congruently rotated objects in EVC

In Experiment 1 (*N* = 35), the occluder was removed during the final scene viewpoint, so that the object reappeared. On 75% of trials, the object reappeared in an orientation that was Congruent with the rotation of the surrounding scene, while on the remaining 25% it was Incongruent (**Figure 1A**). Importantly, the same physical stimuli counted as Congruent or Incongruent depending only on the trial context, avoiding any stimulus-related confounds. We compared participants’ performance in an orthogonal perceptual task (see **Materials and methods** and **Figure 2**), as well as BOLD activity patterns in our two ROIs, evoked by Congruent and Incongruent reappearing objects.

Behaviorally, participants were more accurate on Congruent than Incongruent trials (mean hit rate: 0.64 vs. 0.60, *t*_34_ = 2.99, *p* = 0.005, *d* = 0.67, CI = [0.01, 0.06], **Figure 2B**). This indicates that the rotation of the scene influenced participants’ perceptual processing of the objects.

**Figure 3.**
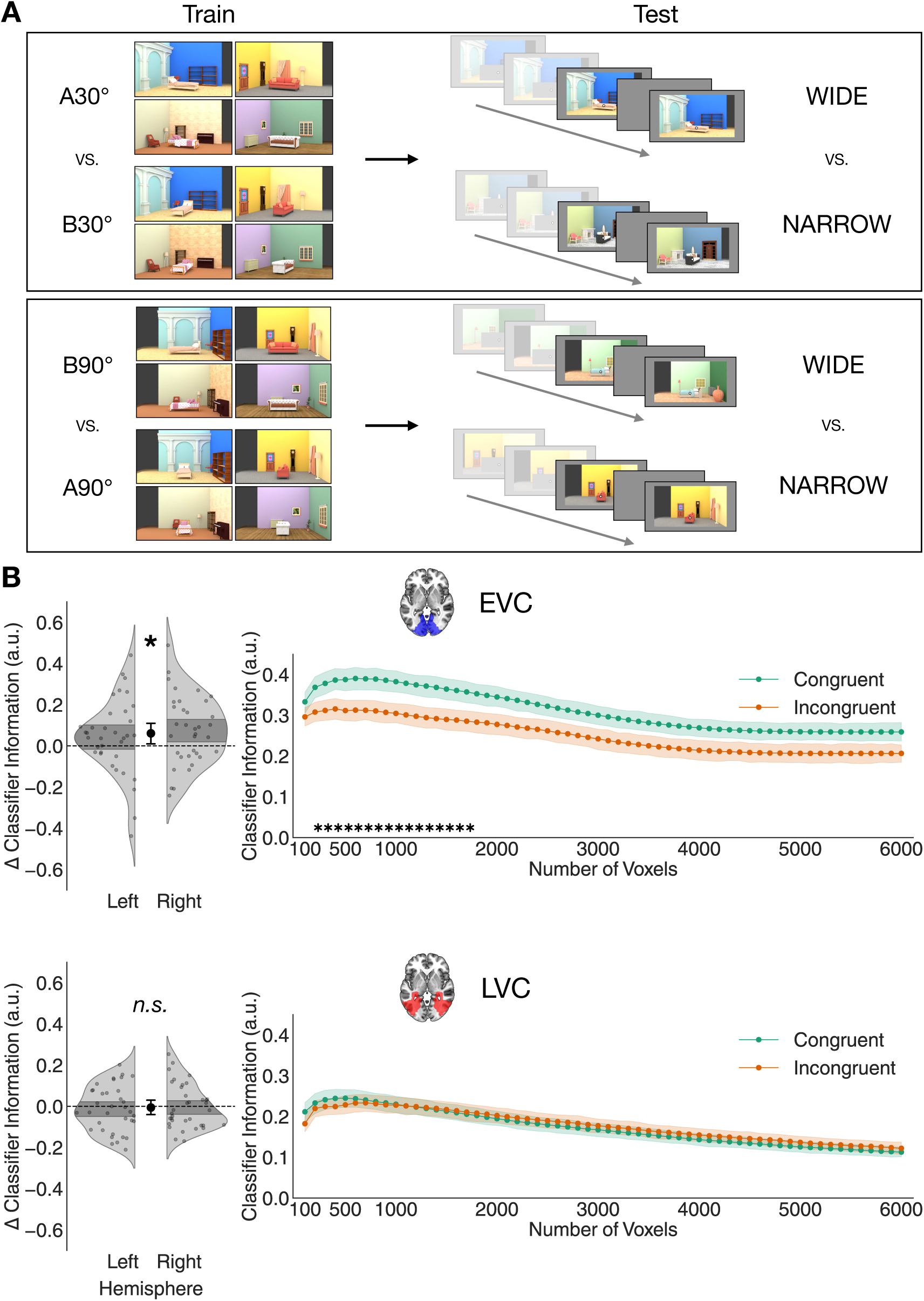
Results of Experiment 1. **(A)** The cross-decoding scheme used in Experiment 1. Linear classifiers were trained to distinguish wide and narrow object views from cortical responses obtained in separate training runs. The stimuli in these runs were the final views shown in the main task runs, but presented without any preceding rotation sequence. To ensure that decoding was driven by the object’s proximal shape, and not by covarying features such as the overall orientation of the scene, separate classifiers were trained to distinguish wide and narrow views with different background orientations (30° and 90°). These were then tested on the (wide versus narrow) object views shown at the end of rotation sequences in the main task runs. Classifier information was then averaged across backgrounds, and compared between Congruent and Incongruent trials. **(B)** Multivariate decoding results of Experiment 1: as the number of voxels to be selected in each ROI (based on the functional localizer) was arbitrary, we varied this number between 100 and 6000 in steps of 100 voxels, creating 60 sub-ROIs with an increasingly liberal inclusion criterion. Classifier information was then averaged across sub-ROIs, and the difference between Congruent and Incongruent was computed for each participant and each hemisphere. This difference is shown on the left side: classifier information was significantly higher for Congruent than Incongruent object views in EVC, indicating that more information about the proximal object shape was present in this ROI. On the other hand, this difference was not found in LVC. The right side shows that these results were consistent across numbers of included voxels, averaged across participants and hemispheres (shaded regions denote SEM across participants). Asterisks denote significance of the difference between Congruent and Incongruent classifier information after applying TFCE (see **Materials and methods** for details). * P < 0.05

To examine the information about Congruent and Incongruent objects in visual cortex, we trained linear classifiers to distinguish the object’s proximal shape (wide versus narrow projection) from BOLD activation patterns. These classifiers were trained on separate training runs, in which all possible final object and scene orientation combinations were shown without the preceding rotation sequence (**Figure 3A**). The purpose of these training runs was to estimate benchmark visual cortical responses to wide versus narrow object orientations, regardless of their contextual (in)congruency. Overall, the proximal shape of the objects could be decoded reliably above chance in both EVC (mean classifier information 0.28, *t*_34_ = 13.85, *P* < 0.001, *d* = 2.39, CI = [0.24, 0.32]) and LVC (mean classifier information 0.11, *t*_34_ = 11.86, *P* < 0.001, *d* = 2.00, CI = [0.14, 0.20]). Thus, information about the object’s appearance was present throughout the visual cortex. Decoding accuracy was significantly higher in EVC than LVC (*t*_34_ = 7.11, *P* < 0.001, *d* = 1.02, CI = [0.08, 0.14]), likely due to the stronger sensitivity of earlier visual areas to changes in an object’s appearance, such as across viewpoints (*22*).

Turning to our central analysis, we found that the shape of Congruent objects could be cross-decoded better than the shape of Incongruent objects in EVC, and this was consistent across a large range of voxel inclusion thresholds (**Figure 3B**, Congruent vs. Incongruent means across voxel numbers: 0.31 vs. 0.25, *t*_34_ = 2.38, *P* = 0.023, *d* = 0.44, CI = [0.01, 0.11]). This result was consistent in both directions of decoding (classifiers trained on training runs and tested on main task runs, or vice versa, **Figure S7**), and when using classification accuracy instead of classifier information (**Figure S4**). It was also consistent when only including the subset of trials in which the last visible object orientation was perfectly matched across small and large scene rotations (**Figure S2**). This indicates that the enhancement of object representations was driven by the scene context, and not by differences in the visible object orientations before occlusion (see **Materials and methods** for more details on this control analysis). On the other hand, no difference in cross-decoding performance between Congruent and Incongruent objects was found in LVC (**Figure 3B**, Congruent vs. Incongruent means across voxel numbers: 0.17 vs. 0.18, *t*_34_ = -0.37, *P* = 0.71, *d* = 0.06, CI = [-0.04, 0.03]). To confirm that the difference between Congruent and Incongruent cross-decoding was stronger in EVC, we ran a within-subject ANOVA with congruency (Congruent, Incongruent) and ROI (EVC, LVC) as factors. This analysis revealed a significant interaction between congruency and ROI (*F*_1,34_ = 12.18, *P* = 0.0014, *η^2^_p_* = 0.26). Congruency with the scene’s rotation, then, enhances the information present in visual cortex about the object’s proximal shape, and this effect appears to be specific to early stages of visual processing.

**Figure 4.**
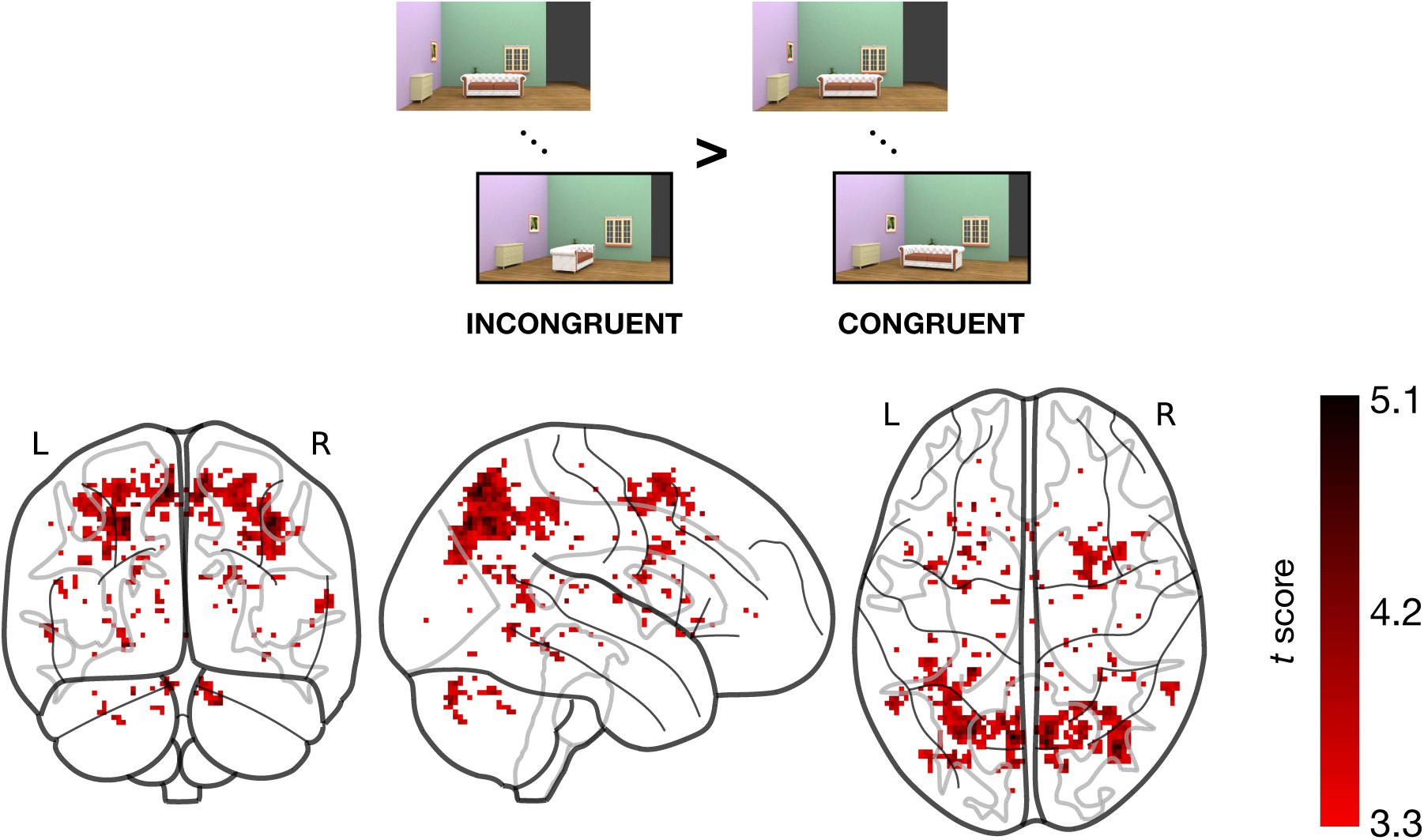
Results of the univariate contrast between Incongruent and Congruent trials. Several clusters responded more strongly to Incongruent trials, while none responded more to Congruent ones. This result suggests that incongruently oriented objects elicited a ‘surprise’ response.

### Incongruent objects elicited a larger univariate response

We next investigated whether the observed enhancement in multivariate decoding was accompanied by an overall higher univariate response. If participants were actively anticipating the appearance of an object that matched their scene-driven expectations, it is possible that attention to Congruent objects would lead to a larger univariate response (*23–25*) . For example, a larger response would be expected if participants were actively maintaining the Congruent object in working memory (*25*) or if attention was captured by the Congruent object (*23*, *24*, *26*). A higher signal-to-noise ratio in conditions with overall higher response amplitudes could then underlie the better multivariate decoding in the Congruent condition. Alternatively, the enhancement of object information in EVC could have occurred in the absence of a higher univariate response, or even with a lower response. This would be more consistent with expectations resulting in a more efficient neural code (*20*, *21*).

In EVC, which showed enhanced decoding for congruent object information, we did not observe any difference in univariate response, independently of the number of voxels included in the analysis (**Figure S3**): Congruent vs. Incongruent means across voxel numbers -1.98 vs. - 1.90, *t*_34_ = -1.05, *P* = 0.302, *d* = 0.04, CI = [-0.22, 0.07]. In fact, the mean activation on Congruent trials was numerically lower. This result indicates that the enhanced multivariate decoding we observed in EVC does not result from an overall larger univariate response.

We next ran a whole-brain univariate contrast, to determine whether any clusters in the brain display a significantly higher response to either Congruent or Incongruent objects. There were no clusters responding more to Congruent than Incongruent objects. Conversely, several clusters responded more to Incongruent than Congruent objects (**Figure 4**, **Table S1**). The most prominent clusters were found in the precuneus, angular gyrus and inferior parietal lobe, areas associated with attentional reorienting and cognitive control. Together, these results indicate that the congruency of objects with the rotation of the scene evoked an overall smaller, not larger, univariate response. This finding is consistent with the idea of congruent object representations being represented more efficiently in the visual cortex (*20*, *21*). Moreover, it reinforces the conclusion of our recent behavioral work (*15*), that scene-driven object predictions are generated automatically rather than as a product of active and voluntary mental operations.

### Multivariate enhancement co-varied with activation in higher-level visual cortex

The contextual enhancement observed in EVC required the integration of information across large regions of the visual field. Moreover, it was based on high-level scene information. For these reasons, it most likely involved computations occurring in higher-level visual or associative areas. The previously reported enhanced decoding of degraded objects embedded in scenes, for example, is driven by feedback from scene-selective cortex (*10*, *27*, *14*). To reveal which brain regions were involved in the enhancement we observed, we ran an information-activation coupling analysis (*28*). This analysis determines whether the univariate activation of particular voxels co-varies, across timepoints after stimulus onset, with the accuracy of multivariate decoding in a seed region, in our case EVC. In particular, we tested whether this coupling was stronger in the Congruent than the Incongruent condition (see **Materials and methods** for details). Locations in the brain that are more strongly coupled with the decoding accuracy in the seed region on Congruent than Incongruent trials are likely to be involved in the enhancement of Congruent object representations.

**Figure 5.**
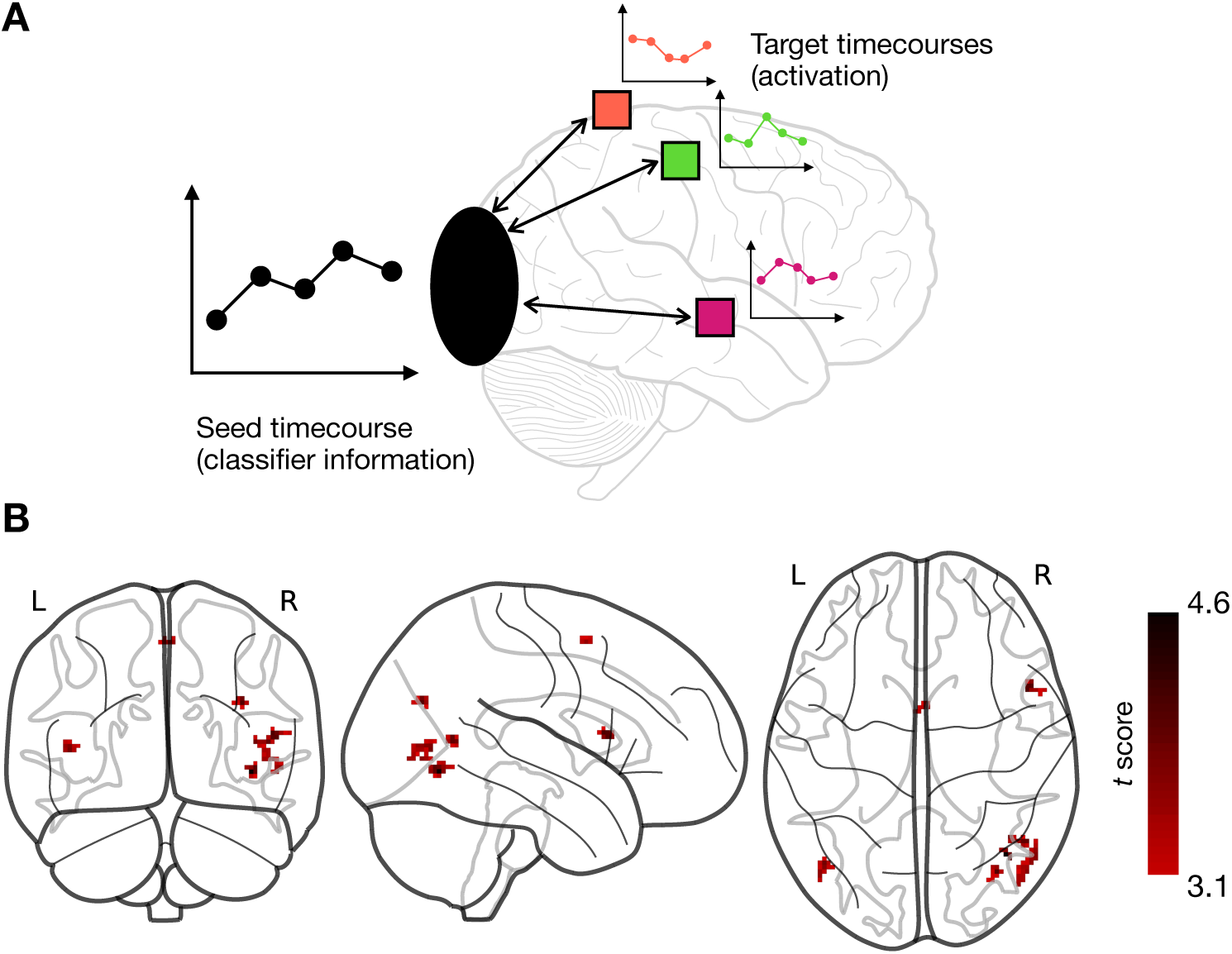
Information-activation coupling analysis. **(A)** Illustration of the information-activation coupling analysis. Given a seed timecourse of multivariate classifier information in an ROI (in our case, EVC) after stimulus onset, and a target timecourse of univariate activation for each voxel in the brain, the per-voxel correlation with the seed timecourse is computed across the whole brain. These correlations are then compared between the Congruent and Incongruent conditions, to reveal voxels that are more strongly coupled with multivariate information in EVC in the Congruent condition. **(B)** Results of the one-sided univariate contrast between the correlation maps for Congruent and Incongruent trials. Several clusters were found that were significantly more coupled on Congruent than Incongruent trials, corresponding to higher-level visual cortex, parietal, premotor and inferior frontal cortex (see text for details).

We contrasted the coupling for Congruent and Incongruent conditions across the whole brain, as we did not have strong prior hypotheses about which regions might be the source of dynamic scene-driven predictions. This analysis revealed several clusters showing greater coupling for Congruent than Incongruent objects (Figure 5 and **Table S2**). We used the Neurosynth platform (*29*) to search for the terms most strongly associated with the peak coordinates of these clusters, based on meta-analysis maps. This search (see **Table S3**) revealed that two of the clusters were associated with visual motion and motion-sensitive area V5/MT (most associated terms: “visual motion”, “v5”, “motion”, “mt”) as well as with object processing (“fusiform”, “objects”, “object”). Other clusters were most strongly associated with the inferior frontal gyrus and premotor cortex (“inferior frontal”, “premotor”, “imitation”, “handed”), as well as with parietal cortex and spatial cognition (“spatial”, “parietal occipital”, “visuo”, “navigation”). These results suggest that the object predictions we observed involved the interaction between EVC and higher-level visual areas related to motion and object processing, as well as the inferior frontal gyrus, premotor and parietal cortices, which were previously implicated in coordinate transformations and mental rotation (*30*, *31*).

**Figure 6.**
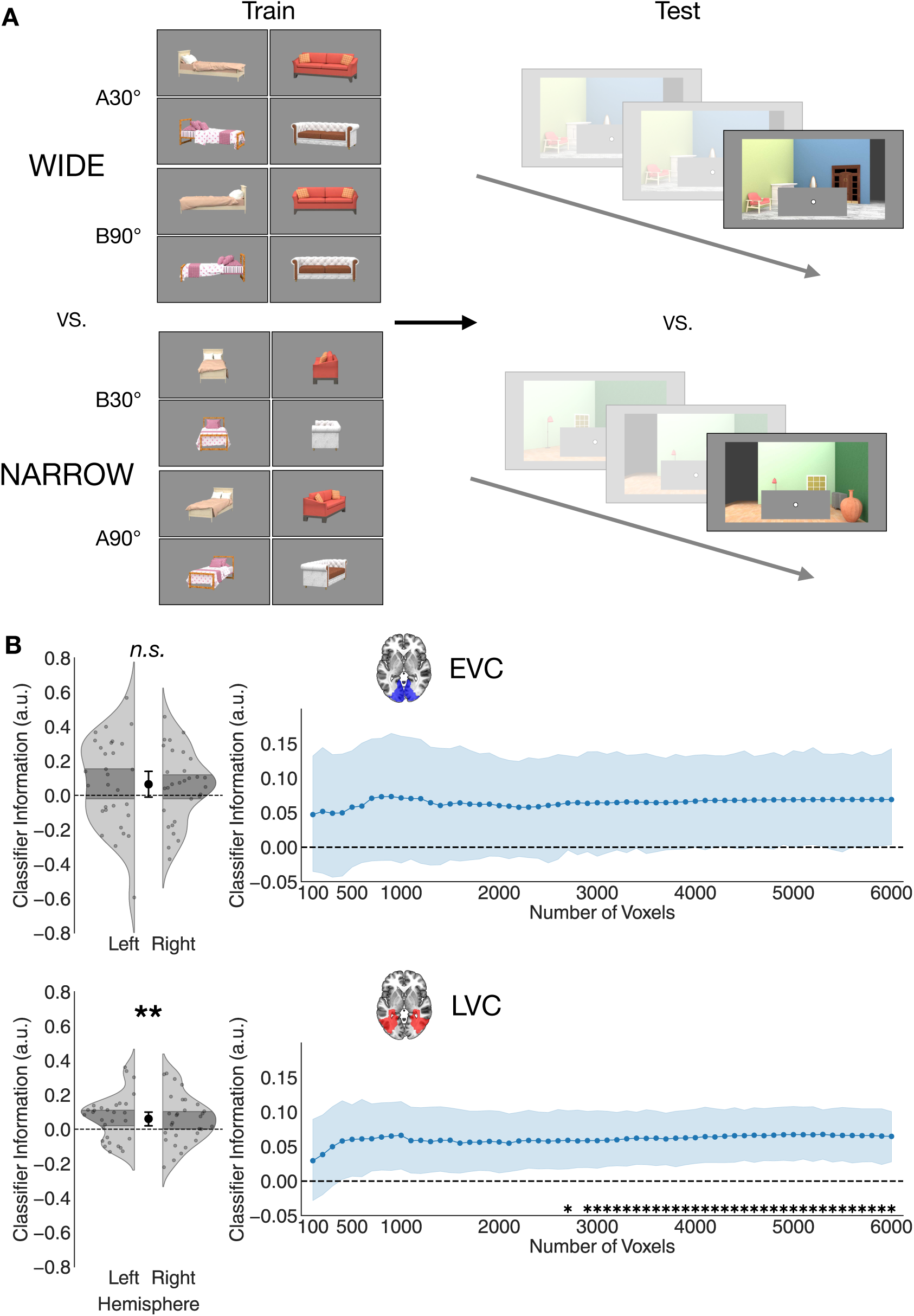
Results of Experiment 2. **(A)** Multivariate cross-decoding scheme used in Experiment 2. Linear classifiers were trained to distinguish wide and narrow object views from BOLD activity in training runs. In these runs, objects were shown without any background. Object views were grouped in the ‘wide’ and ‘narrow’ categories based on their proximal shape, independently of their orientation relative to the scene and the viewer. Thus, the ‘wide’ category included both A30° and B90°, and the ‘narrow’ category included both B30° and A90°. The classifiers thus trained were then tested on the final period of trials in the main task runs, in which the scene had completed its rotation, and the object was still occluded. The goal was to determine whether an expectation of the occluded object’s rotated view was present in visual cortex despite the object not being visible. **(B)** Results of the multivariate decoding analysis of Experiment 2. Varying the number of voxels included in the analysis, we found that the expected proximal object shape could be reliably decoded above chance in visual cortex. In particular, classifier information was positive regardless of the number of included voxels in LVC, although the difference between LVC and EVC was not significant, indicating that information about object shape was present throughout visual cortex (see text for details). Left: distribution of classifier information, averaged across voxel numbers, for each participant and hemisphere. Right: classifier information, averaged across hemispheres, for each number of included voxels. Shaded regions indicate SEM across participants, and asterisks indicate significance after TFCE. * P < 0.05, ** P < 0.01

### Scene rotation updated object representations in the absence of visual input

Experiment 1 showed that scene-driven predictions about occluded objects enhance visual cortical object representations. This may indicate that a representation of the predicted object shape is pre-activated in visual cortex, based on changes in scene viewpoint. Alternatively, it may reflect a reactive signal that distinguishes ‘congruent’ from ‘incongruent’ scenes (i.e., devoid of any visual content) and modulates visual cortex activity. Previous work has found that expectations based on environmental regularities, beyond modulating visually evoked activity, can also drive the inference of occluded parts of visual scenes (*5*, *6*, *8*), and elicit visual predictions in the absence of sensory input (*17*, *18*). In Experiment 2 (*N* = 30), we therefore set out to directly investigate whether changes in scene viewpoint pre-emptively evoke a visual representation of the predicted object, while the object was fully invisible (i.e., during occlusion). The experimental design was largely consistent with that of Experiment 1, except that in Experiment 2, the central object remained occluded until the end of the trial. This made it possible to directly examine the internal representation of the object. Moreover, in this experiment participants did not have to actively perform a visual discrimination task on the object, providing a strong test for the automaticity of scene-driven object updating. To ensure that they still paid attention to the stimulus sequence, the object reappeared on 12.5% of trials. When the object did reappear, it was always oriented congruently. At the end of each run, participants had to report the number of reappearances within the run. The data from these catch trials was excluded from all subsequent analyses.

As in Experiment 1, we trained linear classifiers on separate runs to discriminate the proximal shape of objects (wide or narrow, **Figure 6A**). In this case, the classifiers were trained on BOLD responses to visually presented objects without any background, and tested on BOLD responses to scenes with occluded objects, thus cross-decoding from visually evoked to purely top-down responses. We analyzed the same ROIs as in Experiment 1, EVC and LVC, again testing for robustness across varying numbers of included voxels.

We found that the object’s proximal shape could be decoded above chance in visual cortex, consistently across a wide range of voxel numbers (**Figure 6B**). In LVC, decoding was reliably above chance (mean classifier information across sub-ROIs: 0.061 ± 0.002 SEM, *t*_29_ = 2.96, *P* = 0.006, *d* = 0.54, CI = [0.02, 0.1]). This result was consistent without any participant exclusions, despite noisier data (**Figure S6**), when using classification accuracy instead of classifier information as a measure of decoding (**Figure S4**), and across both decoding directions (**Figure S7**). On the other hand, decoding was not significantly above chance in EVC: mean classifier information across sub-ROIs was 0.064 ± 0.004 SEM, *t*_29_ = 1.72, *P* = 0.095, *d* = 0.31, CI = [-0.01, 0.14]. However, a paired t-test comparing classifier information across the two ROIs revealed no significant difference between them (*t*_29_ = 0.12, *P* = 0.902, *d* = 0.02, CI = [-0.06, 0.07]), suggesting that information about the occluded object’s orientation was not strongly confined to any specific region. This result shows that viewpoint changes in a scene can elicit expectations of object appearance in visual cortex, even when the objects are fully invisible.

## Discussion

Vision in complex real-world environments often requires inferring the properties of temporarily invisible objects (*32*). The ability of the human visual cortex to predict incomplete visual scenes has been studied extensively (*5*, *6*, *33*), but it is still an open question how this ability can generalize to predictions based on complex regularities of the environment, such as its 3D geometry. Here, in two neuroimaging studies, we show that activity patterns in visual cortex reflect predictions of the appearance of an occluded object across viewpoint changes in a 3D scene.

In Experiment 1, we found that the proximal shape of objects (a wide versus narrow projection on the 2D image plane) in a rotating scene was decoded better when the objects emerged from occlusion in an orientation that was congruent with the new scene viewpoint, compared to incongruent objects. This multivariate enhancement was accompanied by an overall reduced amount of (i.e., univariate) brain activation, consistent with an effect of expectation on visual representations (*20*, *21*). Interestingly, in Experiment 2, we found that these predictions of object appearance from scene viewpoint could elicit visual representations in the absence of visual input. We found that the proximal shape of the rotated object (i.e., a wide versus narrow projection) could be decoded from visual cortex activity, even when the object remained fully occluded. These results show that temporarily invisible objects can evoke a visual representation, as informed by the surrounding (visible) scene context. To do so, the visual cortex capitalizes on the predictable way in which objects in the real world rotate coherently with the surrounding scene. Such seamless integration of visible and invisible information can be extremely useful in tracking objects across periods of invisibility, as is often required in daily life (*32*).

Across our two experiments, as well as in previous behavioral studies (*15*, *16*), we found multiple converging lines of evidence suggesting that scene-driven predictions occur automatically: (1) They can affect behavioral performance in an orthogonal task (Experiment 1, and previous behavioral studies (*15*, *16*)), and can be reliably decoded in the absence of any explicit task (Experiment 2). Thus, they do not seem to be driven by task requirements. (2) In Experiment 1, we found that incongruent objects elicited a larger univariate response than congruent ones. If participants were actively maintaining an attentional template of the predicted object, we would expect objects matching this template to be boosted, evoking a higher response (*23*, *24*, *26*, *25*). (3) In two previous behavioral studies (*15*, *16*) with large participant samples (N = 152, 151) we found that the effect of scene-driven expectations on task accuracy was entirely uncorrelated with participants’ responses in a questionnaire assessing their behavioral strategy. This provides further evidence that these expectations were not driven by an explicit, voluntary strategy. Relatedly, the effect of scene-driven expectations was more likely to be driven by real-world structural regularities rather than by short-term regularities extracted during the experiment. While in Experiment 1 congruent object orientations were shown on a majority of trials, thus conflating their real-world congruency with their short-term frequency, in previous behavioral studies (*15*, *16*) we found that short-term frequency was not necessary for congruency effects on task performance. Even when the congruent object view was shown on a minority of trials, participants’ predictions still followed real-world (i.e., scene-congruent) expectations. Moreover, in Experiment 2, in which the congruent object orientation was only shown on a small minority of trials (remaining under occlusion on the majority of trials), the predicted orientation could still be decoded above chance, providing further evidence for the automatic and long-term nature of these scene-driven expectations. Overall, the automaticity of these effects and their dependence on real-world regularities, rather than short-term contingencies, supports their potential relevance to real-world predictive vision.

In Experiment 1, the modulatory effect of scene-object congruency on orientation decoding was specific to EVC, suggesting that expectations modulated relatively low-level visual representations. Possible features driving this effect include the differences in retinal size, stimulated area, or edge orientations between wide and narrow views of the objects. The present experiment was not designed to distinguish between the roles of various low-level features, however, leaving this question open for future investigation. Conversely, in Experiment 2, orientation decoding of the occluded object was significant only in LVC. This discrepancy across experiments may reflect that purely top-down generated object predictions are represented relatively more coarsely, providing a ‘scaffold’ to modulate more fine-grained, stimulus-evoked responses in EVC through feedback projections. This interpretation is in line with the results of the coupling analysis of Experiment 1, showing that univariate activity in occipitotemporal cortex (in the proximity of motion-and object-selective regions) was more strongly coupled with congruent than incongruent orientation decoding in EVC. This dissociation has been observed in previous studies as well: EVC has been implicated in a wide variety of cognitive, but visually-based, processes, including mental imagery (*34*), working memory (*35*), mental rotation (*36*, *37*), tracking of occluded objects (*38*), and intuitive physics (*39*, *40*). All these cognitive operations share a fundamentally spatial nature: they require maintaining or manipulating visual information at a specific location in (retinotopic) space. On the other hand, very similar processes that are less spatially specific seem to involve LVC instead. For example, mental imagery only involves EVC when the location and scale of the stimulus to be imagined is clearly specified, and it involves LVC otherwise (*34*). Similarly, the scene-driven modulation of a visible object’s perceived size in the Ponzo illusion (*41*, *42*) occurs in EVC (*43*, *44*), while the size of an object’s search template, a top-down signal without a specific position in the scene, is observed in LVC (*19*). Consistently with these prior findings, in our study, the difference in tasks between Experiments 1 and 2 potentially also contributed to the discrepancy in the involved processing regions. The task of Experiment 1 involved a precise perceptual comparison of two subtly different stimuli, which likely requires the fine-grained spatial resolution of EVC. Experiment 2, on the other hand, featured a much less visually precise task (detecting the reappearance of an object from occlusion), which may not require the fine-grained precision or retinotopic organization typical of EVC.

The present work focuses on investigating the *content* of expectations based on scene context: a prediction of the object’s proximal shape. Future work should investigate the *format* of the representations that make these expectations possible. One possibility is that the scene is represented as a structural description in allocentric 3D coordinates (*45–47*), and then translated back to retinotopic coordinates, leading to the egocentric 2D shape predictions we observed in our study. This kind of explicit coordinate transformation has been proposed to underlie spatial navigation and mental imagery (*48*, *49*). Alternatively, predictions might be represented exclusively in terms of egocentric views, with no involvement of explicit 3D descriptions. Human behavior in spatial navigation tasks, for example, is consistent with scene representations in terms of 2D views (*50–52*). Moreover, recent work has shown that objects’ proximal shape is represented explicitly in several tasks that involve 3D structure, such as mental rotation (*53*, *54*) or searching for objects at different depths in a scene (*19*, *55*). Future studies could shed light on the representations underlying scene-driven predictions, for example by investigating how these predictions are affected by 3D features (such as the angle of rotation) and 2D features (such as egocentric motion patterns), as done in recent work on mental rotation (*53*).

Regardless of whether the representations that participants relied on in our study are based on egocentric views or 3D structure, our results suggest that humans can represent scene-object relations in a sufficiently rich manner to support predictions across changes in viewpoint. This extends a long line of empirical and theoretical work investigating how the internal representation of objects reflects their properties in the external world (*56–58*). This includes the ability to mentally rotate objects (*59*) or to simulate their physical dynamics (*60*). It is possible that these internal representations also incorporate models of how objects interact with their context, including (but not limited to) how objects rotate concurrently with the surrounding scene. One way to efficiently process these kinds of spatial relations in complex scenes is to represent them in a hierarchical manner, linking scenes to the objects they contain, and linking objects to their parts. These kinds of hierarchical representations are extensively used in computer graphics and game engines (*61*, *62*), and artificial intelligence research has addressed the problem of how they can be extracted from unstructured visual input (*63–68*).

Some evidence exists that humans process scenes hierarchically (*69–71*), suggesting that a similar representation might underlie the present results. Alternatively, the link between scenes and objects might be represented in a ‘flat’, non-hierarchical manner, similar to relations between objects (*72*) or social interactions between agents (*73*). To adjudicate between these two alternatives, future studies could test whether the effect of scene rotation on object representations are asymmetric – scenes can rotate objects, but not vice versa, arguing for hierarchical representations, or symmetric, arguing for flat representations.

In conclusion, the current findings show that the visual cortex generates visual predictions derived from the 3D structure of scenes. These findings suggest that previously reported mechanisms for perceptual predictions (e.g., of partially occluded 2D images) generalize to structured and dynamic real-world environments.

## Materials and methods

### Participants

Participants were recruited through the Radboud University participant pool (SONA systems) and received a monetary reimbursement for their participation. They provided informed consent before the experimental session. The study was conducted in accordance with the institutional guidelines of the local ethical committee (CMO region Arnhem-Nijmegen, The Netherlands, Protocol CMO2014/288). For both studies, we aimed to collect a pre-determined sample size of 34, in order to achieve 80% power for detecting a medium-sized (*d* > 0.5) within-subject effect using a two-tailed one-sample or paired *t*-test. In Experiment 1, a total of 35 participants took part in the study (21 females, mean age = 24.1, SD = 4.4). In Experiment 2, a total of 34 participants took part, of which 4 were excluded due to not paying sufficient attention to the stimulus sequences, as measured through our simple task of counting object reappearances. Specifically, at the end of each main task run, participants were asked to report how many times the object had reappeared after the occlusion period. The Pearson’s correlation, across runs, between the true and reported number of object reappearances was used as a measure of each participant’s accuracy. Specifically, participants were excluded when their average correlation was more than 2 inter-quartile ranges away from the first quartile.

The final sample size was then 30 participants (16 females, mean age = 25.2, SD = 8.5). These participants’ responses were positively correlated with the true values (mean *r* = 0.89, minimum = 0.39), with a majority (25/30) having correlations higher than 0.80, as shown in **Figure S5**.

### Apparatus

Participants viewed the stimuli through a mirror mounted on the head coil of the scanner. In Experiment 1, stimuli were presented on a 32-inch BOLDscreen monitor (Cambridge Research) with 1920 x 1080 px resolution and 120 Hz refresh rate. The total viewing distance (eyes from mirror + mirror from screen) was 1206 mm. In Experiment 2, stimuli were presented on an EIKI LC-XL100 projector with 1024 x 768 px resolution and 60 Hz refresh rate, back-projected onto a projection screen (Macada DAP diffuse KBA) attached to the back of the scanner bore. The total viewing distance was 1440 mm. In both experiments, stimuli were presented using Psychtoolbox (*74*) in MATLAB R2017b. Participants provided responses on a HHSC-2x4-C button box.

### Stimuli

In both experiments, the stimuli for the main task and classifier training runs were 20 different indoor scenes (**Figure S1**) modeled in Blender 2.80 and rendered using the Cycles rendering engine for realistic lighting. The scenes all had the same layout (floor, two walls at a right angle and a main object in the center), but contained various additional objects, adjacent to the walls, and different textures on the walls and floors, to increase their perceptual variability. The central object was a couch for half of the scenes, and a bed for the other half. The retinal size of the central objects was approximately the same across scenes. For each scene, a range of viewpoints was rendered, by rotating the entire scene around the vertical axis (out of the image plane) between 0° and 90°, in steps of 5°. A subset of these viewpoints was presented on each trial: the trial always started with the 0° viewpoint, and ended either with 30° or 90°. The three intermediate viewpoints were chosen in the following way: for 30° rotation trials, a fixed sequence (15°, 20°, 25°) of intermediate viewpoints was shown. For 90° rotation trials, three intermediate viewpoints were randomly sampled (without replacement) from the set {15°, 20°, 25°, …, 55°, 60°}, and shown in increasing order. The object was occluded after the first intermediate viewpoint. In Experiment 1, this meant that the last visible object viewpoint could differ between 30° and 90° rotation trials. This did not substantially affect the results, since consistent findings were obtained when using a sub-selection of trials in which the last visible object viewpoint was perfectly matched between rotation conditions (**Figure S2**). Moreover, in Experiment 2 the last visible object viewpoint was fixed at 10° for both rotation conditions, thus avoiding any potential influence of the last visible object orientation on the decoding of the expected object orientations.

In all scenes, the two walls were oriented such that the scene was fully visible from all the viewpoints. The scenes were presented at the center of the screen with a size of 20.53 x 11.64 degrees of visual angle (dva), surrounded by a gray background. The occluder was a gray rectangle (same color as the background) which had the height and width of the largest possible view of the object in that particular scene (average size: 5.50 x 2.86 dva), plus a margin (horizontal: 1.08 dva, vertical: 0.43 dva) to ensure the object was fully covered and its shadow was not visible, which would have provided a cue to its orientation. The fixation dot (radius: 0.1 dva, shown at the center of the central object, 3.24 dva below the center of the screen) was always visible on top of the images.

In Experiment 1, the stimuli for the classifier training runs were the final views of the objects shown in the Main Task runs, with the scene background (**Figure 3A**). In Experiment 2, they were the same objects but without the scene background (**Figure 6A**). The size of the stimuli was the same as in the Main Task runs.

### General procedure

In Experiment 1, before the fMRI scanning session, participants performed a short practice session (40 trials, around 10 minutes duration) to familiarize themselves with the main task of the experiment. During this session, they received feedback on every trial, and they saw their overall accuracy at the end of the session as well. After the practice, they were also instructed about the other tasks they would have to perform in the scanner (one-back task in the Classifier training and Functional Localizer runs). During the five-minute anatomical scan, they practiced the main task again, also with trial-by-trial feedback. In total, participants were in the scanner for 12 functional runs (∼75 minutes). Each functional run began and ended with 15 seconds of fixation.

In Experiment 2, given the less challenging task, there was no practice session. Before entering the scanner, participants were instructed about the main task they were going to perform and were shown example stimuli. They were also told that on some runs they would have to detect repeated images (one-back task in the Classifier training and Functional Localizer runs). During the five-minute anatomical scan, they practiced the main task, receiving feedback. Participants were in the scanner for a total of 13 functional runs (∼70 minutes). One participant included in the final sample (and one excluded participant) only completed 7 main task runs instead of 8.

### Procedure: main task runs

In Experiment 1, participants completed 7 runs of the main task, each consisting of 48 trials (336 trials in total). Within each run, 36 trials (75%) featured the Congruent object orientation at the end of the stimulus sequence and the remaining 12 (25%) the Incongruent orientation. We chose to present Congruent orientations on a majority of trials because our previous behavioral work (*15*) revealed that the behavioral accuracy difference between conditions was highest with this design (although the effect remained present even when the Incongruent trials outnumbered the Congruent trials). By choosing the design in which the effect was strongest, we maximized the power for uncovering the neural correlates of this behavioral effect. Both Congruent and Incongruent trials were equally divided among the 4 possible initial orientation/amount of rotation combinations (A30°, A90°, B30°, B90°).

Crucially, the behavioral task that participants had to perform was fully orthogonal to the congruency manipulation: they did not have to explicitly judge whether the object remained in the same orientation relative to the beginning of the trial, or to explicitly predict its upcoming view after the occlusion period. Participants were told that their task pertains exclusively to the final viewpoint, but were nonetheless instructed to remain attentive during the whole stimulus sequence. Each trial (**Figure 2A**) began with a fixation dot for 500 ms, followed by the initial view of the scene for 2000 ms. The scene then started rotating, in 3 intermediate views, each shown for 500 ms. The object was fully occluded starting from the second of these intermediate views. The final view of the scene, with the object still occluded, was displayed for a randomly jittered time between 1500 and 2000 ms. The object then reappeared and was briefly flashed twice (with the scene background always present) for 50 ms each, with a 100 ms inter-stimulus interval in between. We refer to these two brief presentations of the object as the *probes*. On a given trial, the second probe was rotated clockwise or counterclockwise, with equal probability, relative to the first, and participants’ task was to indicate ‘clockwise’ or ‘counterclockwise’ using the index or middle finger of their right hand, respectively. Participants had a maximum of 1500 ms to respond, after which the experiment would skip to the next trial and the current trial would be counted as missed. The duration of the initial fixation period for the following trial was adjusted to compensate for participants’ response time on the current trial, to ensure that the overall duration of each run was constant. The first probe’s orientation was randomly sampled from a normal distribution centered around the Congruent or Incongruent orientation (depending on the current trial’s condition), with a standard deviation of 1°, to add a small amount of jitter, and then rounded to the nearest integer. The second probe was rotated, clockwise or counterclockwise, relative to the first by an angle that was titrated using a 2-down 1-up staircase, to keep the task difficulty constant across participants. To ensure that the visual stimuli in Congruent and Incongruent trials did not differ, and thus avoid any stimulus-related confounds, a single staircase was used across both Congruency conditions, allowing for accuracy differences between conditions.

Unlike in the practice session, participants did not receive feedback on every trial, to avoid any possible effects on the fMRI response of differing feedback between Congruent and Incongruent conditions. Instead, their overall accuracy within a run was displayed at the end of the run.

In Experiment 2, participants completed 8 runs of the main task (40 trials each) for a total of 320 trials. The stimulus sequence and durations were the same as in Experiment 1. The main difference was that on a majority of trials, the central object was not shown again after the occlusion period. It was shown only on 40/320 trials (12.5%), randomly spread across the 8 runs (between 2 and 10 per run). On these trials, the occluder disappeared, revealing the object in the final orientation (there was no congruency manipulation in this experiment) for 200 ms. To encourage participants to pay attention to the stimulus sequence, at the end of each run they were asked to report on how many trials the object reappeared. An adjustable number (initially set to 0) was shown on screen and participants could increase it using their middle finger or decrease it using their index finger. To confirm their estimate, they used their ring finger. They were then shown both their estimate and the correct number as feedback.

### Procedure: classifier training runs

The purpose of the classifier training runs was to estimate benchmark response patterns to the central objects used in our main task, without the context of the whole rotation sequence. In both experiments, participants completed 3 training runs.

In Experiment 1, the images displayed in the training runs were the final frames of the sequences shown in the main task. They were presented in mini-blocks corresponding to the 4 possible object orientation/scene rotation combinations (A30°, A90°, B30°, B90° – see **Figure 3A**). Each mini-block consisted of 18 images (different scene exemplars, all in the same orientation/rotation combination), with each image presented for 350 ms and followed by a 400 ms blank interval (each mini-block lasted 13.5 s in total). After a series of 4 mini-blocks (54 s), a longer blank interval was shown for 6.75 s. Participants’ task was to press any button whenever the exact same image was repeated twice in a row (one-back task). Each run included 20 mini-blocks (divided into 5 blocks).

In Experiment 2, the objects in the training runs were shown without any scene background (**Figure 6A**). Aside from the absence of a background, the position and size of the stimuli was the same as in the main task runs. Different object exemplars were grouped in mini-blocks by their proximal shape, such that a given mini-block contained exclusively wide or exclusively narrow objects, including different initial orientation and rotation combinations (*wide* mini-blocks included A30° and B90°, *narrow* mini-blocks B30° and A90°). Each mini-block consisted of 9 images (6.75 s in total), each image being shown for 350 ms and followed by a 400 ms blank interval. After a series of 8 mini-blocks (54 s), a longer blank interval was shown for 6.75 s. Participants performed the same one-back task as in Experiment 1.

### Procedure: functional localizer runs

In both experiments, participants completed 2 runs of a functional localizer scan used for ROI voxel selection. The stimuli used in the functional localizer runs in both experiments were the same as those in a well-established functional localization study (*75*). They included images from 4 stimulus categories (objects, scrambled objects, faces and scenes) shown in separate mini-blocks, each lasting 15 s and comprising 20 unique images. Each image was shown for 450 ms and followed by a 300 ms blank. Each localizer run included 16 mini-blocks (divided into 4 blocks, each containing all 4 stimulus categories in varying order). Participants performed the same one-back task as in the classifier training runs. Stimuli were shown with a size of 12 x 12 dva, against a uniform gray background.

### Acquisition and preprocessing of fMRI data

In Experiment 1, fMRI data were collected on a 3T MAGNETOM Skyra MR scanner (Siemens AG, Healthcare Sector, Erlangen, Germany) using a 32-channel head coil. Functional data was acquired using a T2*-weighted gradient EPI sequence, with 6x multiband acceleration factor (TR 1s, TE 35.2 ms, flip angle 60°, 2x2x2 mm isotropic voxels, 66 slices). For the main task runs, 404 images were acquired per run, 333 and 318 images for the classifier training and functional localizer runs, respectively.

In Experiment 2, fMRI data were collected on a 3T MAGNETOM PrismaFit MR scanner (Siemens AG, Healthcare Sector, Erlangen, Germany) using a 32-channel head coil. Functional data was acquired using a T2*-weighted gradient echo EPI sequence, with 6x multiband acceleration factor (TR 1s, TE 34 ms, flip angle 60°, 2x2x2 mm isotropic voxels, 66 slices). For the main task runs, 315 images per run were acquired, and 333 and 318 images for the classifier training and functional localizer runs, respectively.

In both experiments, at the start of the scanning session, a high-resolution T1-weighted anatomical scan was acquired using an MPRAGE sequence (TR 2.3 s, TE 3.03 ms, flip angle 8°, 1x1x1 mm isotropic voxels, 192 sagittal slices, FOV 256 mm). The data was preprocessed using SPM12 (*76*) functions through the Nipype 1.11.0 (*77*) interface in Python. The functional volumes were fieldmap-corrected, spatially realigned, co-registered with the anatomical image, normalized to MNI 152 space using the template provided in SPM, and smoothed with a 3x3x3 mm FWHM Gaussian filter.

### General Linear Model (GLM) estimation

The responses evoked by each of the stimulus types relevant to our analyses were modelled using general linear models (GLMs) in SPM12 (*76*), through the Nipype 1.11.0 (*77*) interface. In both experiments and in all GLM analyses, time series were convolved with the canonical hemodynamic response function (HRF) provided in SPM12.

In Experiment 1, in the main task, the onsets of the final object views were modelled as impulse functions, as they were very rapid visual events. We included regressors for each combination of object orientation and final scene rotation (A30°, A90°, B30°, B90°), separately for the Congruent and Incongruent trials. Since the Congruent condition included 3 times as many trials as the Incongruent condition, estimating beta weights using all trials would have led to a higher signal-to-noise ratio, and consequently a spuriously higher decoding accuracy. To correct this imbalance, we randomly split the 36 Congruent trials within each run into 3 subsets of 12 trials each (thereby matching the number of Incongruent trials). The random splits were determined using a specified seed (different for each subject and run) for reproducibility. Each of the splits was modelled as a separate condition in the GLM, and all subsequent analyses were performed separately on each split, and then averaged. In the classifier training runs, individual mini-blocks were modeled as boxcars. As in the main task runs, we included regressors for each object orientation/scene rotation combination, yielding one beta weight map per condition, per mini-block, per run. For the univariate analysis, we modelled the onsets of the final object views as impulse functions. We only included regressors for the two congruency conditions, Congruent and Incongruent, obtaining two beta weight maps per run.

In Experiment 2, in both the main task and classifier training runs, we only included regressors for the two proximal object shapes (Wide and Narrow), rather than the four separate orientation/rotation combinations. The reason for this was that the objects in the training runs were presented without any background, removing the need to match images by background in the GLM and MVPA analyses (**Figure 6A**, also see **Multivariate pattern analysis**). In the training run mini-blocks, objects were also grouped by their proximal shape regardless of the specific orientation-rotation combination. In the main task runs, the entire period from the onset of the final scene view to its offset was modeled as a boxcar, as we assumed a prediction of the object in its updated orientation would be present throughout this period. Trials in which the object reappeared after the occlusion period were excluded from the analysis. We estimated one beta weight map per run per condition (Wide and Narrow). In the classifier training runs, each mini-block was modeled as a boxcar. We estimated one beta weight map per mini-block, per run, per condition.

In the functional localizer runs of both experiments, mini-blocks belonging to the 4 stimulus categories (objects, scrambled objects, faces and scenes) were modeled as boxcars, yielding one beta weight map per condition per run.

All GLMs included 6 motion parameters and one run-based regressor as nuisance regressors. As participants were performing a one-back task in the classifier training and localizer runs, these runs included an additional nuisance regressor synchronized to participants’ button presses (modeled as impulse functions).

### Regions of interest definition

To select voxels for inclusion in our visual cortex ROIs (in both experiments), we used subject-level t-contrast maps estimated using data from the functional localizers, contrasting stimuli (both objects and scrambled objects) against the fixation baseline. These maps were intersected with an anatomical mask corresponding to Brodmann areas 17 and 18 (corresponding to areas V1 and V2 (*78*)) for EVC, and Brodmann areas 19 and 37 for LVC (*79*). BA19 includes the lateral occipital gyrus and the superior occipital gyrus, while BA37 corresponds to the occipitotemporal cortex and includes the posterior fusiform gyrus and the posterior inferior temporal gyrus. The LVC ROI includes high-level visual regions such as the object-selective LO (lateral occipital) and pFs (posterior fusiform gyrus), as well as the motion-selective hMT+. Each participant’s map, in each hemisphere, was then thresholded to only include the top N most responsive voxels in the stimulus vs. baseline contrast, as measured by the *t*-statistic. The number of selected voxels (N) ranged from 100 to 6000 in steps of 100, creating 60 sub-ROIs per each ROI and hemisphere, with an increasingly liberal voxel inclusion criterion.

### Multivariate pattern analysis

Our cross-decoding analysis consisted of training linear classifiers on benchmark responses (beta weights) to objects devoid of any context (sequence), obtained from the classifier training runs, and testing them on responses to objects appearing at the end of the rotation sequence in Experiment 1 (**Figure 3A**), and on responses to scenes with fully occluded objects in Experiment 2 (**Figure 6A**).

In Experiment 1, in order to decode the stimulus feature of interest – proximal object shape (wide vs. narrow), we separately trained classifiers to discriminate between the A and B object orientations embedded in scenes rotated by 30 or 90 degrees (**Figure 3A**), which corresponds to discriminating conditions A30° and B30°, and A90° and B90°, in such a way as to classify the object’s shape against a matched background. The accuracies of classifiers trained on the two backgrounds were then averaged. The 3 splits of Congruent trials (see **GLM estimation**) were also decoded separately, and accuracy was then averaged across them.

Importantly, the labels of the beta weights corresponding to Incongruent trials in the main task runs corresponded to the object orientation that was *actually* presented on the screen, not the one expected given the context, as our goal was to assess how the same visual stimuli are processed differently depending on the context.

In Experiment 2, as objects were displayed without any background in the classifier training runs, we did not need to implement the background-matched decoding. Additionally, different views that resulted in the same proximal shape were grouped together in the same mini-blocks of the classifier training runs (e.g. A30° and B90° were grouped together as Wide). Classifiers were trained to discriminate between Wide and Narrow objects, and tested on responses to the final views of the scene in the main task runs, where the object was occluded. As the object only reappeared on a small minority of trials, which were excluded from further analyses, these response patterns solely reflected participants’ expectations about the proximal shape of the occluded object.

Besides training on the classifier training runs and testing on main task runs, decoding was also done in the opposite direction (training on main task runs, and testing on classifier training runs) and decoding performance was averaged across directions. This was done because factors unrelated to the task or stimulus, such as different signal-to-noise ratios, can lead to asymmetries between cross-decoding directions (*80*). While our results were consistent in the two decoding directions, they indeed showed different noise levels (**Figure S7**). For all analyses reported in the main text, we thus averaged across directions to obtain a more robust estimate of the stimulus-related information present in multivariate activation patterns. The training and testing datasets were z-scored before decoding.

Multivariate pattern analysis (MVPA) was conducted using linear support vector machines (SVMs) implemented in Scikit-learn (*81*). As a measure of decoding performance, and thus information content in a given brain region, we used the continuous distance from the SVM’s hyperplane (i.e., distance to boundary) rather than discrete classification accuracy.

Continuous measures of the distance between brain activation patterns have been found to be more reliable than discrete ones, likely due to the lossy compression inherent in binary classification outcomes (*82*). Specifically, we used the following continuous measure of decoding performance (which we call *classifier information*):

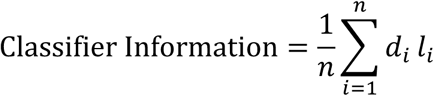

Where *di*’s are the z-scored (across test samples) distances from the hyperplane, *li*’s are the true labels (either -1 or 1) for each sample, and *n* is the number of samples in the test set.

Intuitively, this measure corresponds to the average match between each distance from bound and the corresponding ground-truth label, i.e. the degree to which the distance is positive when the target is positive, and negative when the target is negative. This measure is greater than zero when classification is above chance. The purpose of z-scoring the distances is to remove potential differences between SVMs trained and tested on different data, such as different hemispheres or decoding directions. If the signal-to-noise ratio is higher when training on main task runs, for example, distances in this condition will be higher overall, leading to a disproportionate contribution of this condition when averaging across conditions. Similarly, averaging distances across test samples, rather than summing them, allows us to directly compare classification performance in different conditions, which might have different numbers of samples, and average across them. Specifically, it is necessary for averaging across decoding directions. Classifier information was computed for each sub-ROI within EVC and LVC, in each hemisphere, and each subject. It is important to note that this measure is closely linked to classification accuracy, and all our results were consistent, albeit noisier, when using classification accuracy instead of classifier information (**Figure S4**).

### Significance testing

To statistically test differences in classifier information between conditions (Experiment 1) and absolute amounts of classifier information (Experiment 2), we used two approaches. (1) To avoid making assumptions regarding the appropriate numbers of voxels to include in the analysis for each ROI, we averaged classifier information across numbers of included voxels (sub-ROIs) for each subject and ROI. In Experiment 1, this summary measure was compared between the Congruent and Incongruent conditions with a two-sided paired-sample t-test. In Experiment 2, it was compared against zero with a two-sided one-sample t-test. These statistical tests, as well as the test on behavioral accuracy differences in Experiment 1, were run using Pingouin (*83*). (2) To assess the robustness of (differences in) classifier information across numbers of selected voxels, we used threshold-free cluster enhancement (TFCE) (*84*). TFCE boosts the magnitude of a statistic based on its extent across neighboring samples (in this case, sub-ROIs with similar numbers of voxels), reflecting the assumption that any signal in the data should be smooth across consecutive datapoints. This measure is then compared with a null distribution generated by randomly shifting the signs of each participant’s 1D map (classifier information across sub-ROIs). This null distribution has the same variance and autocorrelation as the original signal. The shuffling procedure was performed 10,000 times. A z-score then expresses how likely each observed TFCE values is, given the TFCE values in the 10,000 permuted (null) data sets, thus implicitly correcting for multiple comparisons. TFCE was computed using the MNE toolbox (*85*).

### Univariate analysis

In Experiment 1, we used a univariate analysis to estimate differences in the overall response elicited by Congruent and Incongruent trials. This was done within the main visual ROIs, as well as across the whole brain. For the within-ROI analysis in visual cortex, we used the same sub-ROIs as in the multivariate analysis, to directly compare the amount of information with the level of activation in the same voxels. We averaged the beta weights across voxels within each sub-ROI (number of selected voxels), separately in EVC and LVC, resulting in one mean beta per condition (Congruent and Incongruent) and participant for each sub-ROI. The averages across sub-ROIs in the Congruent and Incongruent conditions were then compared using a two-sided paired t-test. For the whole-brain analysis, we ran a second-level contrast (one sample two-sided t-test against zero across participants) with α = 0.001 (False Positive Rate corrected), and a cluster threshold of 10 voxels, using the *threshold_stats_img* function in Nilearn(*86*).

### Information-activation coupling analysis

The goal of the information-activation coupling analysis was to reveal regions of the brain in which univariate activation was more strongly correlated with the presence of multivariate information in EVC in Congruent than Incongruent trials. To compute the average timecourses of each voxel in the brain for each condition of interest, we used GLMs with a finite impulse response (FIR) basis function (*87*). We thus obtained, for each condition and run, the BOLD response for 10 time bins (one second each) after stimulus onset (final object appearance). To extract multivariate decoding timeseries, the BOLD activation patterns of EVC in each time bin were fed to an SVM classifier trained to distinguish wide vs. narrow mini-blocks in the training runs. The decoding procedure was the same as in the main multivariate analysis of Experiment 1. This yielded a classifier information score for each time bin for the Congruent and Incongruent conditions. We computed the Pearson’s correlation of these multivariate decoding time series with the time-resolved activation (averaged across runs) in each voxel of the brain, for Congruent and Incongruent conditions. This resulted in two whole-brain maps of correlations for each subject, for the Congruent and Incongruent conditions. To assess robustness to voxel inclusion (for the multivariate decoding in EVC), the whole analysis was repeated for different numbers of included voxels (based on activation in the stimulus vs. baseline contrast, across both hemispheres): 500, 600, 700, 800, 900, and 1000 voxels. The resulting whole-brain maps were averaged. The maps for the Congruent and Incongruent conditions were then compared using a paired-samples t-test, to find voxels that were significantly more correlated with multivariate classification in the Congruent than the Incongruent condition. As we were exclusively interested in clusters that showed more coupling for Congruent than Incongruent trials, we ran a one-sided test. Apart from this, we used the same Nilearn function and parameters as in the univariate analysis of Experiment 1 (see **Univariate analysis**).

## Acknowledgements

We thank Paul Gaalman for MRI scanning assistance.

## Author contributions

Conceptualization: GA, SG, MVP

Methodology: GA, SG, MVP

Software: GA

Formal Analysis: GA

Investigation: GA, SG

Resources: GA

Data curation: GA

Writing—original draft: GA

Writing—review & editing: GA, SG, MVP

Visualization: GA

Supervision: SG, MVP

Funding Acquisition: MVP

## Data and materials availability

The preprocessed fMRI data and beta weights used in all the analyses, the behavioral data, visual stimuli, and the experiment presentation and data analysis code is publicly available at the following link: https://doi.org/10.34973/30xv-q144.

The analysis code is also available at the following repository: https://github.com/GAldegheri/dyncontext.

---

**Figure S1.**
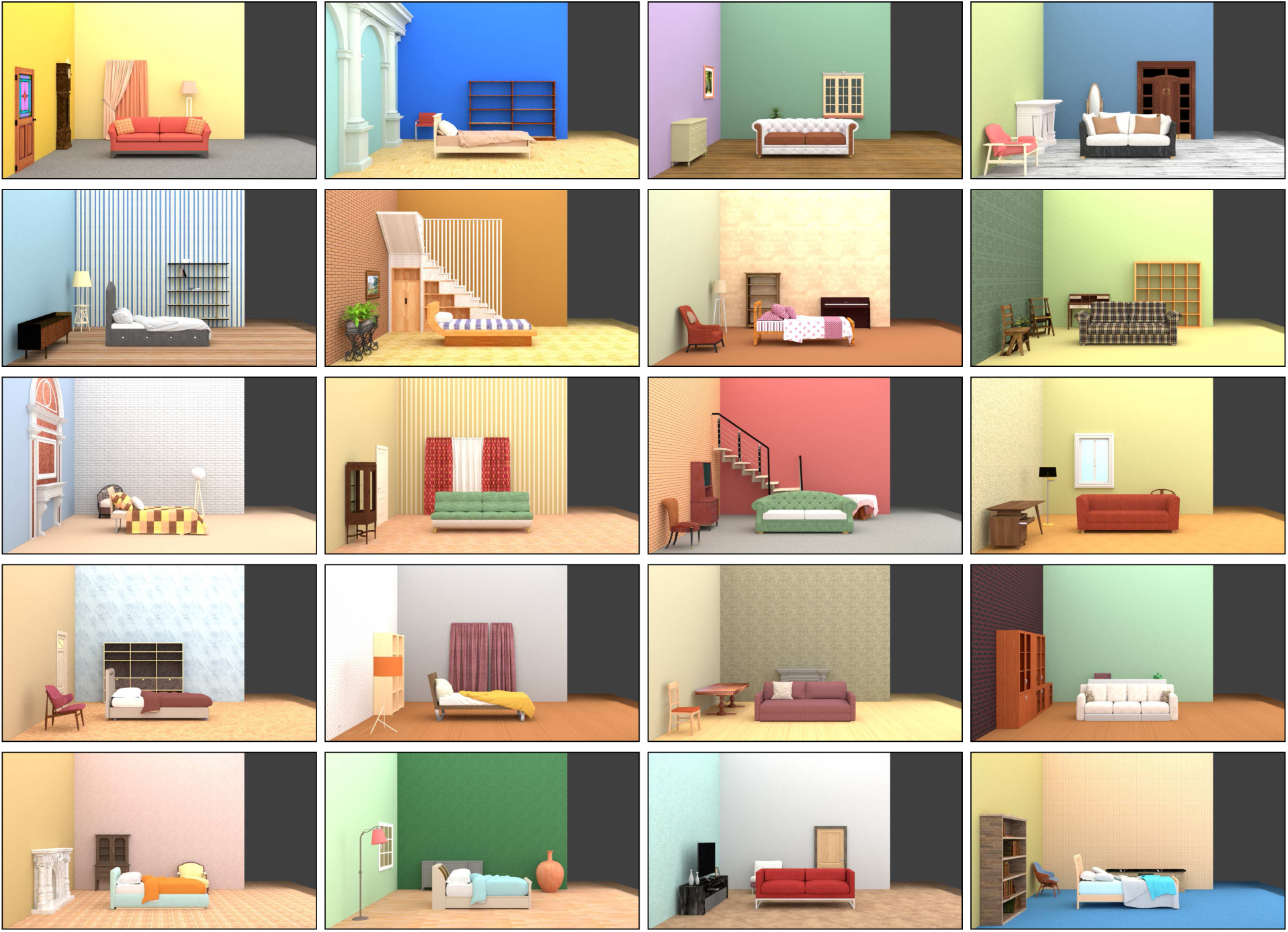
The 20 scene exemplars used in the study.

**Figure S2.**
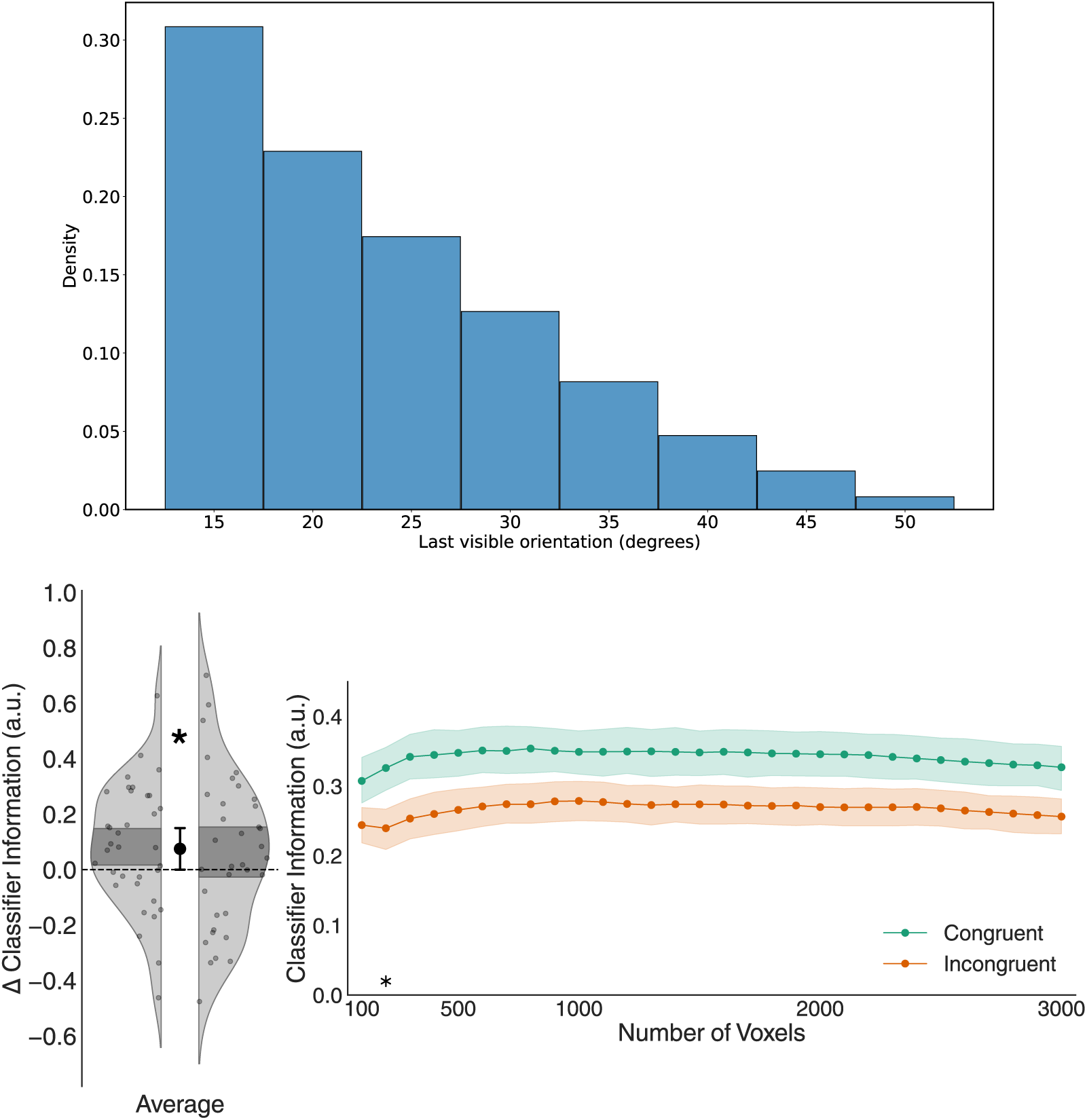
(Top) Distribution of the last visible object orientations, before appearance of the occluder, in the large (i.e., 90°) rotation trials of Experiment 1. In the small (i.e., 30°) rotation trials, this last visible object orientation was always 15°. Thus, in around 30% of the large rotation trials, the last visible orientation of the object was the same as in the small rotation trials, in which case it was impossible to predict the total scene rotation based on this last visible orientation alone. (Bottom) To ensure that the predictive value of the last visible object orientation did not substantially impact the results of Experiment 1, we re-ran the classification analysis of Experiment 1 on EVC, including only the subset of large rotation trials in which the last visible orientation was 15°, thus ensuring that it was perfectly matched (across all trials) with the small rotation condition (for which the last visible orientation was always 15°). As the number of large trials with a last visible orientation of 15° varied randomly from run to run, we had to ensure that the number of trials was balanced between the congruent and incongruent conditions. We thus selected, per each run, the condition (between A90° and B90°, congruent/incongruent) with the smallest number of trials and randomly subsampled trials in the other conditions to match this number. This led to the exclusion of a considerable number of trials. For brevity, this analysis was only run in one decoding direction (training to main task runs), and only selecting subsets of up to 3000 voxels. Despite the substantial reduction in statistical power resulting from the lower number of included trials, this reanalysis of the results of Experiment 1 in EVC replicated the original analysis reported in the main manuscript. * p < 0.05

**Figure S3.**
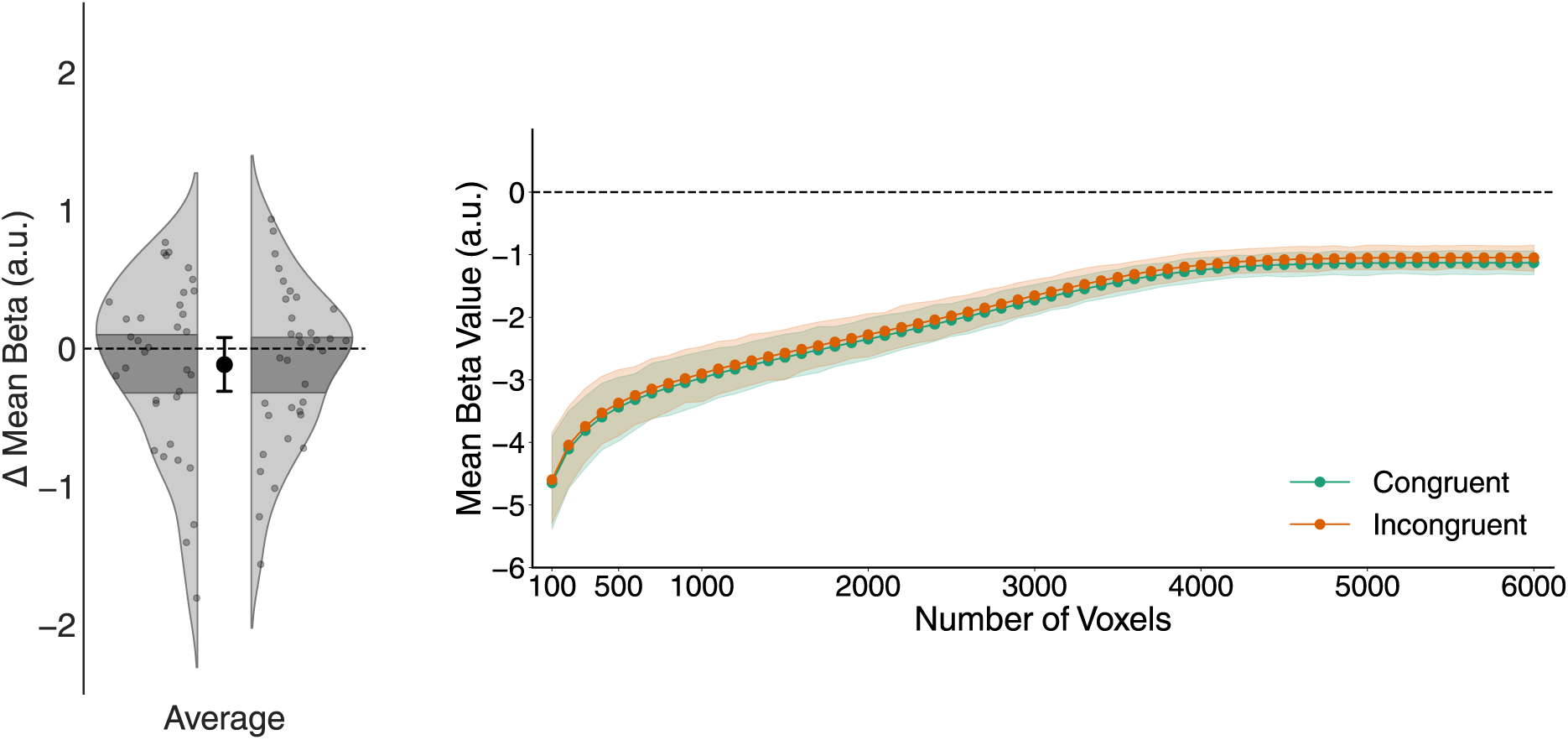
Univariate activation (mean beta value) in EVC for the difference between Congruent and Incongruent trials (left panel) and for Congruent and Incongruent trials separately, across numbers of included voxels (right panel). Univariate activation did not differ in EVC between Congruent and Incongruent trials (and was numerically even higher for Incongruent than Congruent trials), indicating that the increased decodability of object information in Congruent trials did not derive from an increase in overall activation. See Figure 3B in the main text for the corresponding multivariate results.

**Figure S4.**
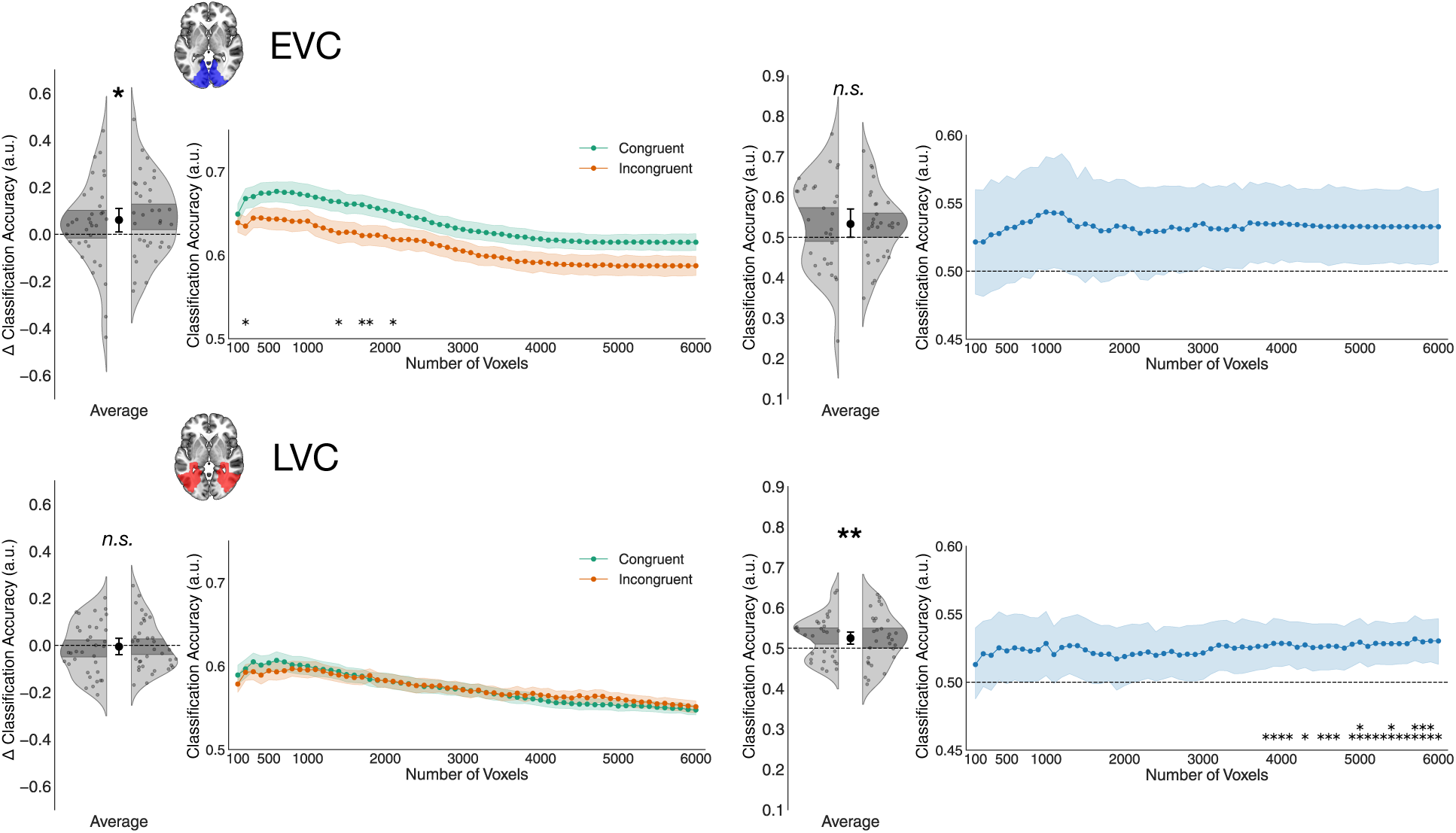
Multivariate decoding results of Experiment 1 (left) and Experiment 2 (right) when using classification accuracy as a measure of information rather than classifier information. See Figures 3B **& 6B** in the main text for the corresponding plots using classifier information.

**Figure S5.**
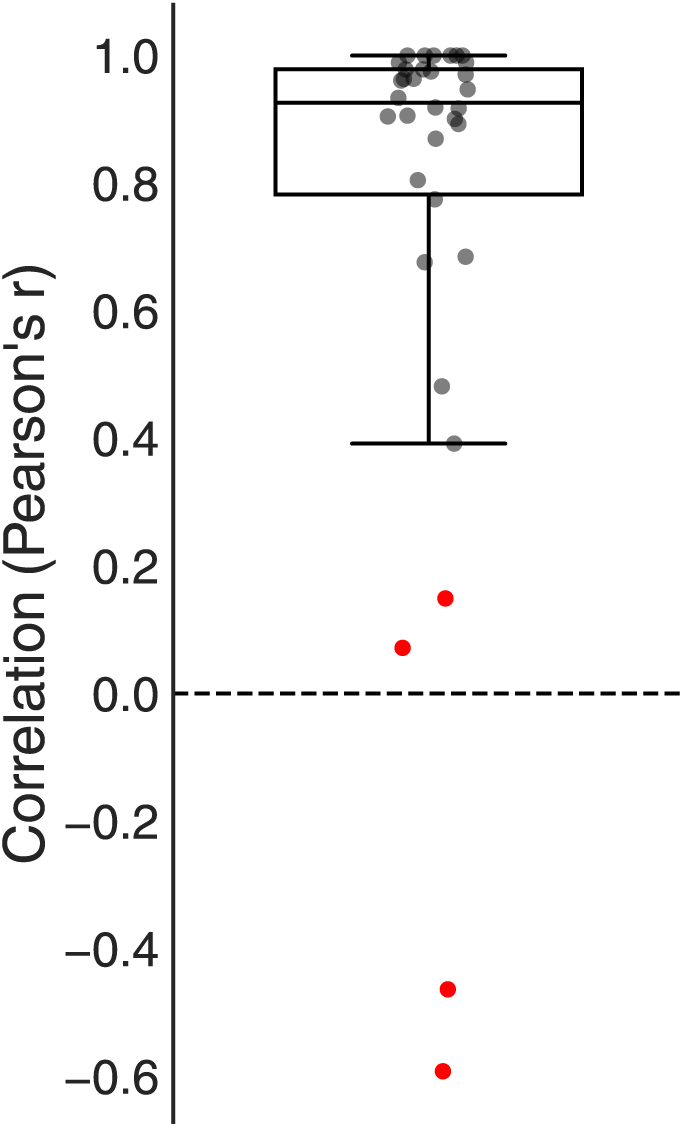
Accuracy in the simple recall task of Experiment 2 for each participant, measured as the Pearson’s correlation between participants’ estimates and the true number of object reappearances. Points highlighted in red indicates outliers (participants who were more than two inter-quartile ranges away from the first quartile), which were excluded from the analysis. The boxplot indicates first, second (median) and third quartile, and the whiskers are drawn until the farthest point within two inter-quartile ranges from the first quartile, or the minimum among the included participants.

**Figure S6.**
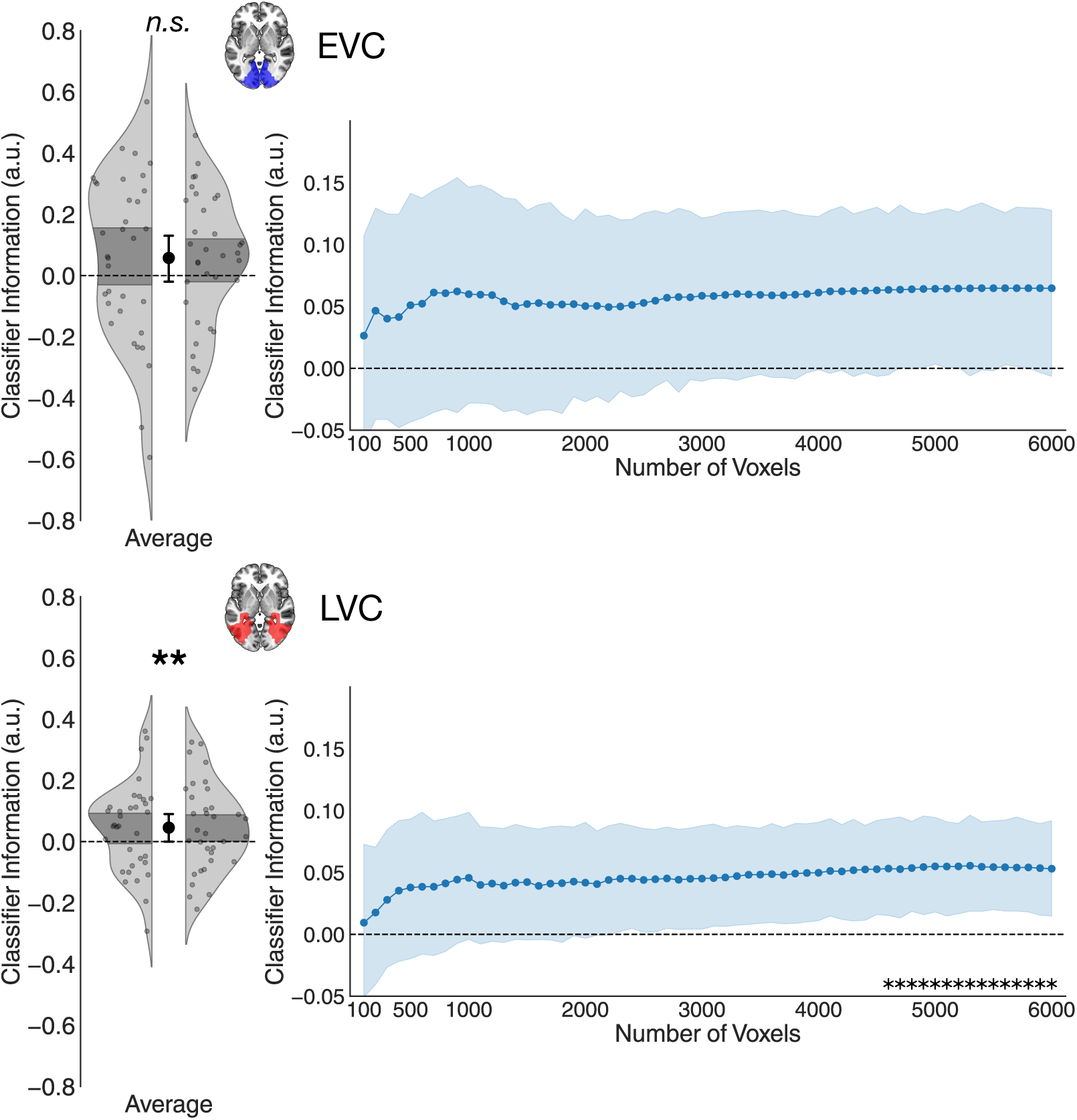
Results of Experiment 2 without any participant exclusions. See Figure 6B in the main text for the corresponding results with participant exclusions.

**Figure S7.**
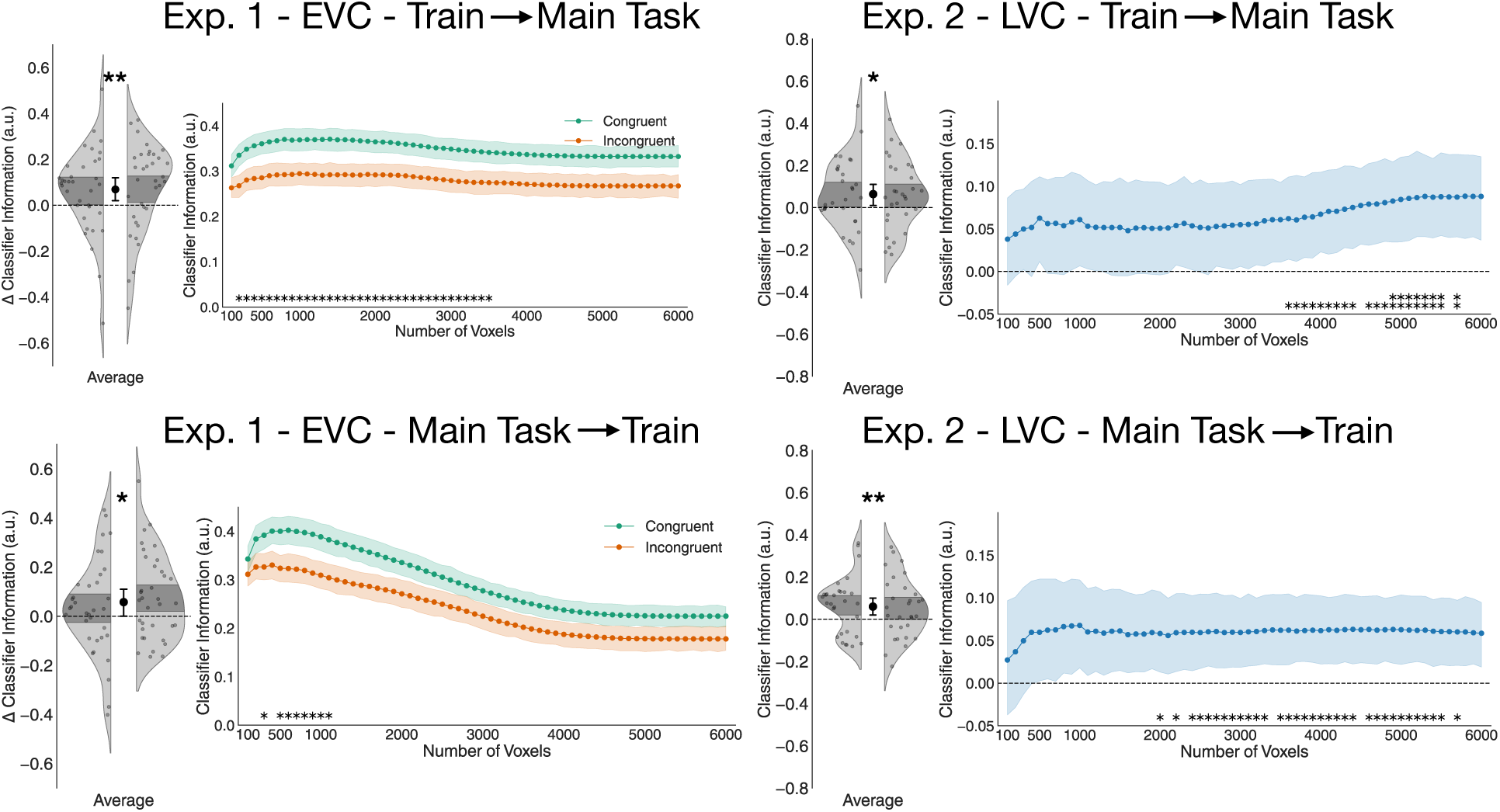
Results for EVC in Experiment 1, and LVC in Experiment 2, separated by decoding direction. The results do not differ in the two directions, although the noise levels are different.

**Table S1.** Clusters showing a significantly stronger response for Incongruent relative to Congruent trials in Experiment 1. No clusters were found that showed a significantly stronger response to Congruent than to Incongruent trials.

| Cluster ID | X | Y | Z | Peak Stat | Cluster Size (mm3) |
| --- | --- | --- | --- | --- | --- |
| 1 | -6.0 | -68.0 | 56.0 | 5.064 | 768 |
| 1a | -10.0 | -62.0 | 52.0 | 4.354 |  |
| 1b | -6.0 | -68.0 | 62.0 | 3.949 |  |
| 1c | -14.0 | -66.0 | 62.0 | 3.601 |  |
| 2 | 36.0 | -72.0 | 42.0 | 4.804 | 1392 |
| 2a | 34.0 | -56.0 | 50.0 | 4.189 |  |
| 2b | 40.0 | -64.0 | 40.0 | 4.092 |  |
| 2c | 34.0 | -58.0 | 42.0 | 3.987 |  |
| 3 | -24.0 | -66.0 | 44.0 | 4.767 | 816 |
| 4 | 8.0 | -70.0 | 54.0 | 4.728 | 840 |
| 4a | 22.0 | -66.0 | 54.0 | 4.136 |  |
| 5 | -22.0 | -40.0 | -8.0 | 4.590 | 96 |
| 6 | 28.0 | -56.0 | 36.0 | 4.388 | 80 |
| 7 | 16.0 | -80.0 | -32.0 | 4.231 | 88 |
| 8 | -26.0 | -6.0 | 52.0 | 4.207 | 104 |
| 9 | 26.0 | -4.0 | 50.0 | 4.185 | 640 |
| 9a | 26.0 | 6.0 | 60.0 | 4.003 |  |
| 9b | 22.0 | 12.0 | 54.0 | 3.819 |  |
| 10 | -26.0 | 8.0 | 58.0 | 4.164 | 144 |
| 11 | -34.0 | -54.0 | 46.0 | 4.147 | 184 |
| 12 | -4.0 | -78.0 | -24.0 | 4.132 | 88 |
| 13 | -30.0 | -76.0 | 32.0 | 4.076 | 192 |
| 14 | -56.0 | -54.0 | 0.0 | 4.042 | 120 |
| 15 | -26.0 | -60.0 | 54.0 | 4.025 | 216 |
| 16 | 20.0 | -56.0 | 24.0 | 4.001 | 120 |
| 17 | -38.0 | -44.0 | 50.0 | 3.941 | 440 |
| 17a | -40.0 | -36.0 | 48.0 | 3.932 |  |
| 18 | 38.0 | 10.0 | 34.0 | 3.841 | 120 |
| 19 | 60.0 | -50.0 | 12.0 | 3.792 | 192 |

**Table S2.** Clusters showing a significantly higher correlation on Congruent vs. Incongruent trials with multivariate decoding time courses in EVC (information-activation coupling analysis) in Experiment 1. This analysis revealed clusters in bilateral higher-level visual cortex, and in parietal, premotor and inferior frontal cortex, that were implicated in the enhancement of object information in EVC.

| Cluster ID | X | Y | Z | Peak Stat | Cluster Size (mm3) |
| --- | --- | --- | --- | --- | --- |
| 1 | 40.0 | -64.0 | -4.0 | 4.559 | 96 |
| 2 | 50.0 | 12.0 | 14.0 | 4.077 | 128 |
| 3 | 36.0 | -72.0 | 28.0 | 4.050 | 96 |
| 4 | 48.0 | -58.0 | 10.0 | 3.934 | 128 |
| 5 | -44.0 | -68.0 | 6.0 | 3.842 | 168 |
| 6 | 54.0 | -64.0 | -2.0 | 3.734 | 136 |
| 7 | 50.0 | -72.0 | 2.0 | 3.698 | 80 |
| 8 | 4.0 | 4.0 | 56.0 | 3.607 | 80 |
| 9 | 46.0 | -78.0 | 2.0 | 3.551 | 80 |

